# Disentangling shape and size in a population of unusually large Threespine Stickleback (*Gasterosteus aculeatus*) from Vancouver Island, British Columbia

**DOI:** 10.64898/2026.04.01.715936

**Authors:** Sara Perry, Kevin K. Duclos, Heather A. Jamniczky

## Abstract

Sarita Lake, British Columbia houses a distinctive population of threespine stickleback (*Gastrosteus aculeatus* L.) with a phenotype characterized by unusually large individuals relative to nearby conspecifics. We tested the hypothesis that members of this population are not isometrically larger but rather exhibit variation in allometric trajectories that reflect changes in developmental timing impacting the developmental-genetic architecture of the phenotype. We used 3D geometric morphometrics to characterize the size and shape of skulls, pectoral girdles and pelvic girdles from a sample of individuals from nearby freshwater and marine populations and compare them to a sample from Sarita Lake. We showed that individuals from the Sarita Lake population are larger in each body region compared to most other populations examined. Further, these individuals have dorsally expanded skulls and relatively robust pelvic armour. We also showed that the relationship between size and shape is differently structured among body regions and is heavily influenced by non-uniform sexually-mediated variation across populations sampled. Our results reflect complex underlying developmental trajectories, and we suggest that the ‘large’ phenotype observed may be driven by fecundity selection on female size in combination with a limnetic trophic niche and relatively increased predation pressure in Sarita Lake.

## Introduction

The threespine stickleback (*Gasterosteus aculeatus* L.) is a small teleost species with robust populations in coastal marine, brackish and freshwater habitats throughout the Northern Hemisphere (Bell and Foster, 1994). The evolutionary history of the threespine stickleback is characterized by rapid diversification as freshwater environments are repeatedly colonized, producing distinctive freshwater and marine populations (Bell and Foster, 1994, Morris et al., 2018). The broad geographic and ecological distribution created by this historical pattern has generated tremendous phenotypic diversity, making the threespine stickleback an invaluable model organism for evolutionary biologists (Bell and Foster, 1994, McKinnon and Rundle, 2002, Reid et al., 2021).

Habitat-specific phenotypic variation has been extensively documented in threespine stickleback as populations adapt to local conditions (Willacker et al., 2010, Webster et al., 2011). Indeed, stickleback inhabiting neighbouring river, salt marsh, estuary, and ditch environments within a single drainage basin exhibit significant morphological variation among sites (Webster et al., 2011). Additional studies have linked these morphological differences to differences in muscle function and swimming performance, further supporting an ecological explanation for this variation (Seebacher et al., 2016). Furthermore, habitat-specific variation in stickleback has consistently been found to be a combination of both genetic influences and phenotypic plasticity in response to environmental factors (Spoljaric and Reimchen, 2011, Wund et al., 2012, McGuigan et al., 2010, Kozak et al., 2025). Phenotypic changes in stickleback can thus take place rapidly in response to novel environmental conditions. For example, anadromous stickleback re-introduced to Loberg Lake, Alaska, where stickleback had been exterminated, exhibited dramatic phenotypic divergence from the ancestral anadromous form within only two years, from 1990 to 1992. Within 17 years, between 1992 and 2009, the introduced stickleback population came to closely resemble the original, extinct Loberg Lake population (Aguirre and Bell, 2012).

Many studies on phenotypic variation in stickleback have focused on parallel evolutionary changes in the skeletal phenotype during freshwater colonization, where freshwater stickleback exhibit a dramatic reduction in the number of lateral armour plates (Bell and Foster, 1994, McKinnon and Rundle, 2002, Reid et al., 2021). This emphasis on the marine-freshwater transition and lateral plates has left gaps in understanding other components of phenotypic variation in stickleback and how developmental constraints may shape phenotypic transitions.

One such developmental constraint is found in allometry, which refers to biological scaling relationships modulating size-related trait variation (Klingenberg, 2016). Allometry is a crucial component of development that can be altered by changes in developmental timing across ontogeny (Alberch et al., 1979). Allometry directly shapes the adult phenotype, and may also constrain variation among populations regardless of genetic or environmental influences, which themselves both facilitate and constrain phenotypic variation along axes that are often, but not always, aligned (Schluter, 2000). Allometric variation in stickleback has been observed alongside phenotypic variation (Aguirre and Bell, 2012, McGuigan et al., 2010, Spoljaric and Reimchen, 2011, Wund et al., 2012, Kimmel et al., 2008, Taugbol et al., 2020). For example, stickleback reared in different controlled environments not only display different adult phenotypes, resulting in distinct ecotypes, but also modified allometric trajectories (Wund et al., 2012). In the case of the stickleback reintroduced to Loberg Lake, allometry was shown to make a small but significant contribution to the eventual phenotypic divergence from the source population (Aguirre and Bell, 2012). In another example, changes in allometric trajectories have been found to occur within populations in response to changing environments, and subsequent transplant experiments demonstrated that the transplanted stickleback population exhibited a distinct shift from allometric to isometric growth in only five years (McGuigan et al., 2010). Opercle development in stickleback has been shown to be strongly allometric, with significant differences in allometric trajectory between anadromous and lacustrine populations (Kimmel et al., 2008).

Sarita Lake is located in the Sarita River watershed on the western coast of Vancouver Island, British Columbia, Canada, and is home to a population of threespine stickleback. The Sarita Lake stickleback exhibits a distinctive phenotype characterized by unusually large size, reported as a mean fork length of 66.4 mm and a mean mass of 2.18g, 33% longer and more than double the mass of stickleback from the nearby Frederick Lake, also part of the Sarita River watershed, which have a mean fork length of 49.7 mm and a mean mass of 0.88g (Martel and Lauzon-Guay, 2005). Known as “gigantism”, unusually large body size is a trait with significant ecological and evolutionary implications that is often produced, at least in part, by the modification of allometric trajectories in isolated populations (e.g. Lefebvre et al., 2022, Hennekam et al., 2020). Cases of gigantism are rare in threespine stickleback but have been documented in Kunk Lake, AK and lakes in Haida Gwaii, BC (Bell, 1984, Reimchen, 1988, Reimchen, 1991, Moodie, 1972, Moodie and Reimchen, 1976).

Whether the Sarita Lake sticklebacks are truly “giant” is debatable. Whereas a mean fork length over 60 millimetres places the Sarita stickleback at the upper end of the usual size range for this species (approximately 30 to 60 mm), the Kunk Lake and Haida Gwaii “giant” populations mentioned above had standard lengths ranging from 66.5 mm up to 115 mm (Bell, 1984, Reimchen, 1988, Reimchen, 1991). Some Sarita Lake individuals do reach sizes well within the giant range, with the largest individual recorded having a fork length of 84.3 mm (Martel and Lauzon-Guay, 2005). Ultimately, the Sarita Lake population seems to occupy an intermediate point between “large normal” and “small giant” but appears anecdotally much larger than most other stickleback populations known from Southern British Columbia. Beyond studies of lateral plate and gill raker polymorphism and asymmetry in the Haida Gwaii stickleback mentioned above, the phenotypes and development of abnormally large stickleback, including those from Sarita Lake, have not been characterized in detail.

Growth, as a developmental process, is well understood to be among the strongest integrators, producing strongly correlated variation among body parts that reflects underlying developmental and genetic integration (Zelditch, 1988, Cheverud, 1996, Klingenberg, 2016, Hallgrimsson et al., 2009). Here, we asked whether and how the anecdotally larger phenotype of the Sarita Lake stickleback population is in fact larger than the phenotype of nearby freshwater populations. We compared the size and shape of skulls, pectoral girdles and pelvic girdles from a sample of individuals from geographically proximate freshwater and marine populations to a sample from Sarita Lake, to test the hypothesis that size variation among populations is not the result of isometric change but rather results from variation among allometric trajectories that reflect changes in developmental timing and growth patterns within the species.

## Material and Methods

### Specimens

Specimens from five populations of threespine stickleback from British Columbia (BC), Canada were examined in this study (Table 1) including three freshwater localities: Sarita Lake (n=55), Hotel Lake (n = 47) and Klein Lake (n = 44); and two marine localities: Bargain Bay Lagoon (n = 49) and Hospital Bay Lagoon (n = 45). No new specimens were collected for this work; rather, existing specimens from a previous study (Schutz et al., 2022) were repurposed for use here. Briefly, adult fish (standard length ≥35 mm, (Baker et al., 2015) were collected using minnow traps from Hotel and Klein Lakes and from Bargain Bay and Hospital Bay Lagoons in 2015 and 2016. Fish were collected from Sarita Lake (48°54′32′′ N, 124°53′26′′ W, perimeter 8.5km, area 126.0 ha, mean depth 14.6 m, maximum depth 28.7 m) in 2015 and 2016. Specimens were euthanized in the field using an overdose of Eugenol (Sigma-Aldrich, Oakville, ON). Fin clips were taken and stored in 70% ethanol, and specimens were then fixed flat in 10% neutral buffered formalin for 24 hours before being stored in 70% ethanol. DNA was extracted from fin clips using a DNEasy Blood and Tissue kit (Qiagen). Sex was determined by genotyping the isocitrate dehydrogenase locus following a standard protocol (Peichel et al., 2004). All sampling and tissue processing were conducted in accordance with the standards of the Canadian Council of Animal Care, and all activities were approved by the Life and Environmental Sciences Animal Care Committee at the University of Calgary (AUPs BI09R-41, AC12 0057, AC13-0040, AC16-0059, and with annual permits approved by the Huu-ay-aht First Nation and the Bamfield Marine Sciences Centre), as described previously.

**Table 1.** Dataset used in this study. Grey shaded rows indicate marine populations.

| Population | Female | Male | Total |
| --- | --- | --- | --- |
| Bargain Bay Lagoon | 23 | 26 | 49 |
| Hospital Bay Lagoon | 14 | 26 | 40 |
| Hotel Lake | 9 | 37 | 46 |
| Klein Lake | 28 | 16 | 44 |
| Sarita Lake | 26 | 29 | 55 |
| Total | 100 | 134 | 234 |

### Models and Landmarking

Three-dimensional reconstruction and surface model generation, as well as landmarking protocols are described in detail elsewhere (Schutz et al., 2022). Briefly, micro-computed tomography instruments (Scanco uCT35 (Scanco Medical AG) or Skyscan 1173 High Energy MicroCT (Bruker)) were used to image all specimens in a standardized position at 20 μm resolution (70 KvP, 114 μm). Raw scans were reconstructed in three-dimensions using either Amira v. 5.4 (ThermoFisher Scientific EM Solutions) or 3DSlicer v. 5.0.3 for Sarita Lake (Fedorov et al., 2012). Landmarks were collected from the right side of the skull, pectoral and pelvic girdles of reconstructed models according to the scheme presented in Figure 1. This landmarking scheme is modified from Schutz et al. 2022, and includes 35 landmarks on the skull, 7 landmarks on the pectoral girdle, and 8 landmarks on the pelvic girdle. Landmarks were collected using either Amira v. 5.4 (ThermoFisher Scientific EM Solutions) or Stratovan Checkpoint v. 2022.12.16 for Sarita Lake (Stratovan Corporation). Landmarks for all study populations were then exported for further analysis. Preliminary data quality analyses were conducted to ensure that no artifacts were introduced using different reconstruction or landmarking software. Landmarks were divided into three sets: skull, pectoral girdle and pelvic girdle, to investigate regionalization within the stickleback skeleton.

**Figure 1.**
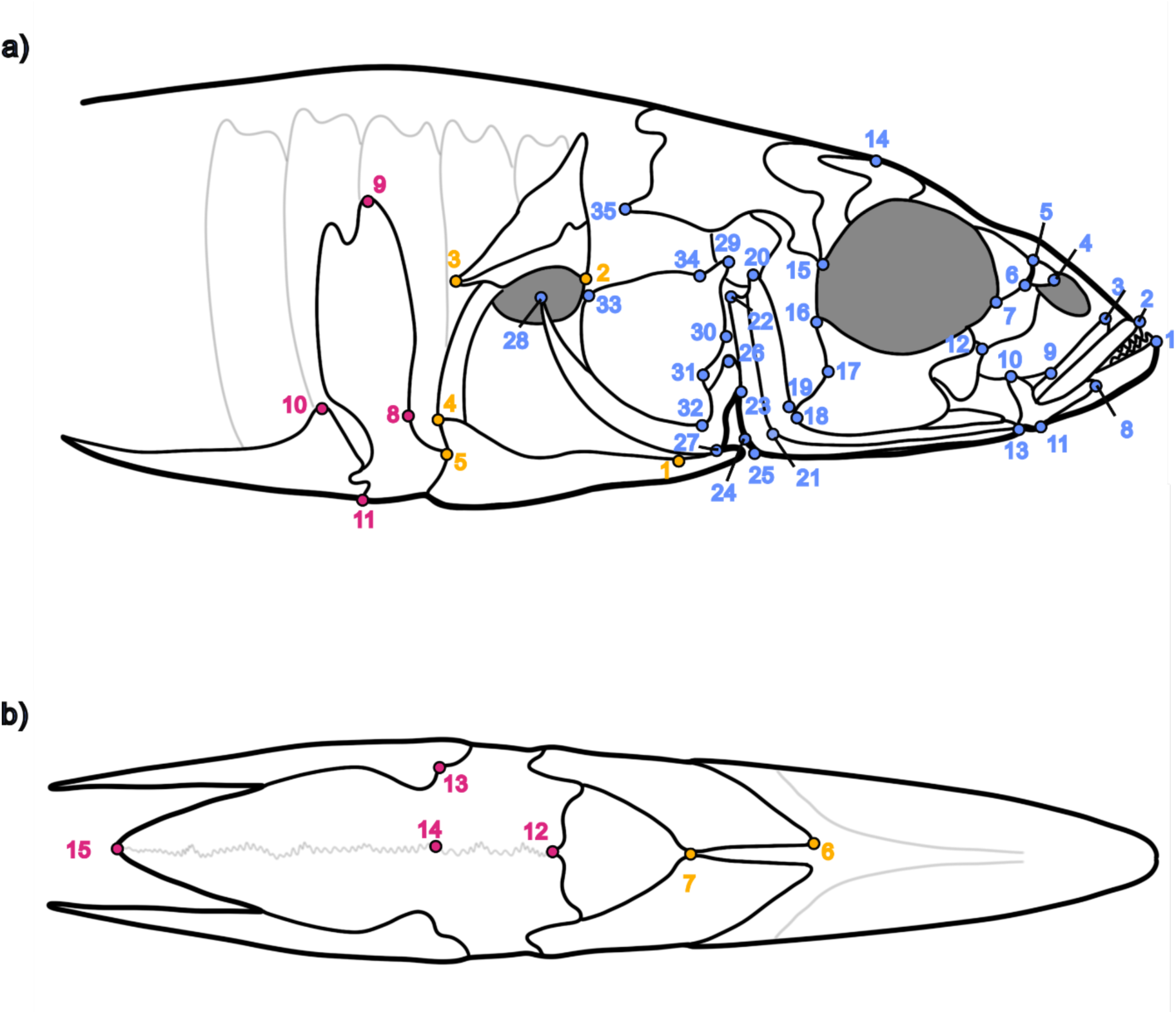
Landmarking scheme for the skull and the pectoral and pelvic girdles in threespine stickleback. The anterior skeleton is illustrated in right lateral view (a) and from a ventral view (b). Skull landmarks (in blue) are as follows: 1, anterior tip of dentary, 2, anterior tip of premaxilla, 3, anterior tip of maxilla, 4, anterior corner of nasal ventrolateral process, 5, dorsal corner of nasal-lateral ethmoid suture, 6, dorsal maximum of lacrimal, 7, lacrimal-prefrontal suture on orbital, 8, anterior tip of articular, 9, ventral maximum of lacrimal, 10, tip of the dorsal process of the articular, 11, ventral-most process of the articular, 12, proximal suture between lacrimal-second orbital, 13, anterior tip of preopercle, 14, dorsal-most extent of supraorbital, 15, ventral-most tip of sphenotic, 16, dorsal-most tip of third suborbital, 17, posterior minimum of third suborbital, 18, ventral-most tip of third suborbital, 19, anterior minimum of preopercle, 20, anterior dorsal-most tip of preopercle, 21, posterior maximum of the first ridge of the preopercle, 22, posterior dorsal-most tip of preopercle, 23, dorsal-most tip of interopercle (note, the structure is not visible on the illustration), 24, ventral maximum of the second ridge of the preopercle, 25, ventral-most tip of interopercle, 26, dorsal-most tip of subopercle, 27, ventral maximum of subopercle, 28, posterior tip of subopercle, 29, dorsal-most tip of opercle, 30, anterior maximum of opercle, 31, anterior minimum of opercle, 32, ventral-most tip of opercle, 33, posterodorsal tip of opercle, 34, opercular hinge angle, 35, posterior tip of pterotic. Pectoral (yellow) and pelvic (pink) girdle landmarks are numbered together: 1, anterior junction between ectocoracoid and coracoid at the caudal-most projection of the coracoid foramen, 2, Anterior-most maximum of the inferior edge curvature of the cleithrum, 3, caudal-most extension of the cleithrum, 4, posterior dorsal-most extension of ectocoracoid, 5, posterior extension of ectocoracoid, 6, anterior tip of ectocoracoid, 7, posterior-most point of the ventral plate of the ectocoracoid, 8, minimum of the anterior curvature of the anterior process of pelvic plate at junction with the base of the ascending branch of the pelvic plate, 9, dorsal most tip of ascending branch of pelvic plate, 10, dorsal most edge of the pelvic spine, 11, ventral most intersection between pelvic spine and ascending branch of the pelvic plate, 12, medial edge of anterior-most point of the anterior process of the pelvic plate, 13, intersection between ventral point of pelvic spine and anterior process of the pelvic plate, 14, medial most point of junction between the anterior process and posterior processes of the pelvic plate at trochlear joint, 15, posterior tip of posterior process of the pelvic plate. This landmarking scheme is modified from Schutz et al. 2022. (Note: Pelvic landmark #16 from Schutz et al. was removed from the analysis due to broken spine tips in our dataset).

### Statistical Analyses

Analyses were conducted independently for the three body regions, which were first aligned separately using a Generalized Procrustes Transformation (GPA) to remove the effects of rotation, translation and scale (Dryden and Mardia, 1998), and then analysed as described below using R v. 4.3.3 (R Core Team, 2025) running in RStudio v. 2024. 12.1 (Posit Software, 2025). Statistical analysis were conducted using geomorph v. 4.0.10 (Adams et al., 2025, Baken et al., 2021) and RRPP v. 2.1.2 (Collyer and Adams, 2024).

Analysis of Variance using residual randomization (RRPP-ANOVA; Collyer and Adams, 2018), using 10,000 permutations, was used to assess the presence of statistically significant differences in the size of each morphological region, represented here by the natural log of the centroid size (logCsize) for the relevant landmark set (Klingenberg, 2016). Pairwise comparisons as implemented in the geomorph ‘pairwise’ function (Collyer et al., 2015) were conducted to identify significant differences in distances among least squares mean centroid sizes.

Principal Components Analysis (PCA) was conducted on each landmark set following GPA to visualize major axes of variation in the dataset. High-dimensional Procrustes RRPP-ANOVA (Randomization of Residuals in a Permutation Procedure; (Collyer et al., 2015) with 10,000 permutations was used to examine shape variation in each landmark set following GPA. Log Csize was included in these models as a covariate, with population, sex, and interaction terms initially included as factors. Backward selection was used to reduce these models where appropriate and the simplest appropriate model was selected. Pairwise comparisons, using the ‘pairwise’ function (Collyer et al., 2015), were conducted to identify significant differences in distances among least squares mean shapes between populations and sexes within populations. Fitted values from these models were then subjected to PCA and these scores were plotted against logCsize to provide a graphical assessment of allometric variation in the sample.

## Results

### Size

Skull centroid size varied significantly among populations and sexually mediated variation was present (Figure 2a, Table 2). Sarita skulls were among the largest in the sample, and Klein Lake among the smallest. Pairwise comparisons (Supplementary Table 1) indicated that the significant differences observed were driven by differences between Sarita Lake female skulls and those of the other freshwater populations, and that the Klein Lake population exhibited sexually-mediated variation in this trait, with males having larger skulls (p < 0.05).

**Figure 2.**
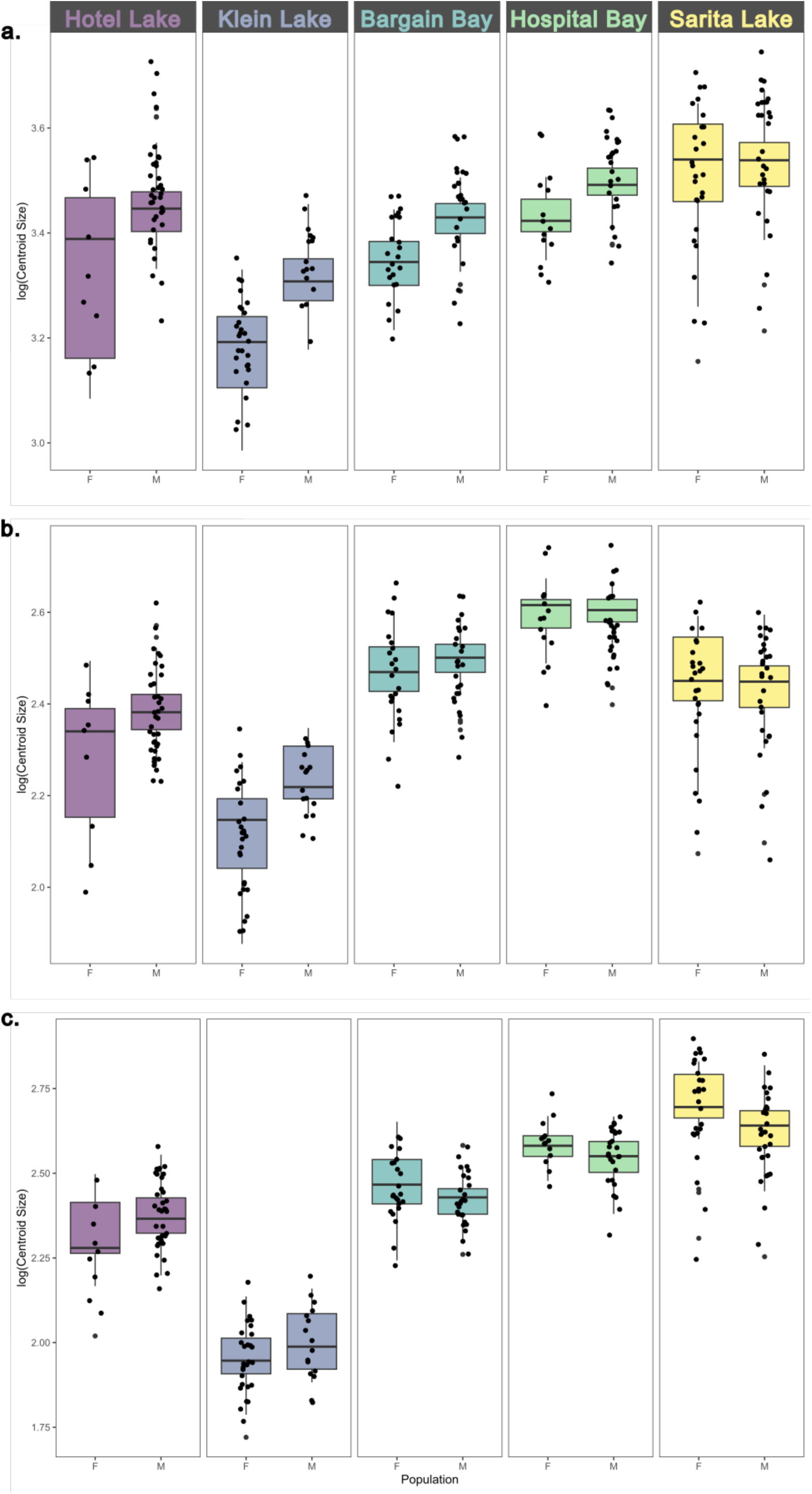
Centroid size variation across the three sampled body regions. a) Skull, b) Pectoral girdle, c) Pelvic girdle. F, female; M, Male. Significant pairwise comparisons (P<0.05) are, for the skull: Hotel Lake F – Sarita Lake F; Klein Lake F – Sarita Lake F; Klein Lake F – Klein Lake M; and for the pectoral girdle: Hotel Lake F – Hospital Bay F, Sarita Lake F, Hotel Lake M; Klein Lake F – Hospital Bay F, Sarita Lake F, Hotel Lake M, Klein Lake M; Hospital Bay F – Sarita Lake M. All pairwise comparisons were significant for the pelvic girdle, with the exception of Hotel Lake – Bargain Bay Lagoon; Hospital Bay Lagoon – Sarita Lake. Note there was no significant sexually mediated variation in the pelvic girdle dataset (see Table 3). Pairwise comparisons are shown in Supplementary Tables 1, 2 and 3).

**Table 2.** RRPP-ANOVA results comparing skull centroid size across all populations examined. See Supplementary Table 1 for significant pairwise comparisons. Abbreviations: Df = degrees of freedom; F = F statistic; MS = mean square; p = p value (10,000 permutations); Rsq = R squared; SS = sum of squares; Z = exect size.

|  | Df | SS | MS | Rsq | F | Z | p |
| --- | --- | --- | --- | --- | --- | --- | --- |
| Population | 4 | 2.2945 | 0.5736 | 0.5179 | 75.9921 | 12.3555 | 0.0001 |
| Sex | 1 | 0.3100 | 0.3100 | 0.0697 | 41.0701 | 4.3166 | 0.0001 |
| Population:Sex | 4 | 0.1353 | 0.0338 | 0.0305 | 4.4824 | 2.8656 | 0.0019 |
| Residuals | 224 | 1.6908 | 0.007 | 0.381 |  |  |  |
| Total | 233 | 4.4307 |  |  |  |  |  |

**Table 3.**
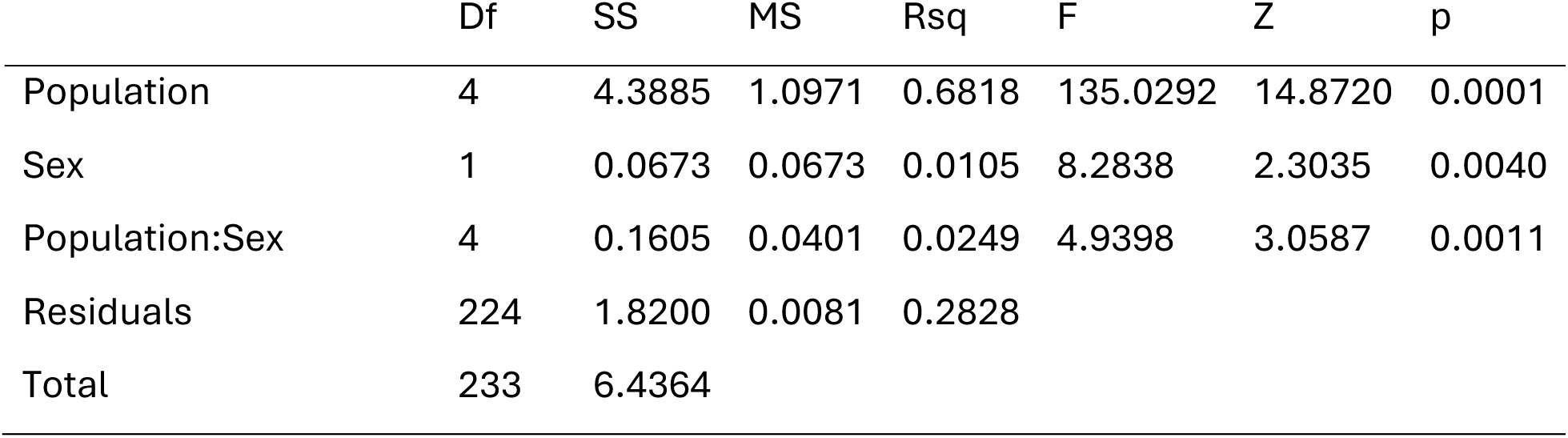
RRPP-ANOVA results comparing pectoral girdle centroid size across all populations examined. See Supplementary Table 2 for significant pairwise comparisons. Abbreviations: Df = degrees of freedom; F = F statistic; MS = mean square; p = p value (10,000 permutations); Rsq = R squared; SS = sum of squares; Z = effect size.

Pectoral girdle centroid size varied significantly among populations and sexually-mediated variation was present (Figure 2b, Table 3), with Hospital Bay pectoral girdles among the largest and Klein Lake among the smallest. Pairwise comparisons (Supplementary Table 2) indicated that the significant differences observed were largely driven by differences between Klein Lake girdles and those of other populations (p < 0.05), and that the Klein Lake population again exhibited sexually mediated variation in this trait, with males having larger pectoral girdles (p < 0.05). Sarita Lake pectoral girdles were generally larger than those of nearby freshwater populations, but smaller than those of marine comparators.

Pelvic girdle size varied significantly among populations, but this body region did not exhibit any sexually-mediated variation (Figure 2c, Table 4). Klein Lake pelvic girdles were significantly smaller than those of all other populations (p = 0.0001), and the Sarita Lake pelvic girdles were significantly larger than those of all other populations except Hospital Bay (p = 0.0001, Supplementary Table 3).

**Table 4.**
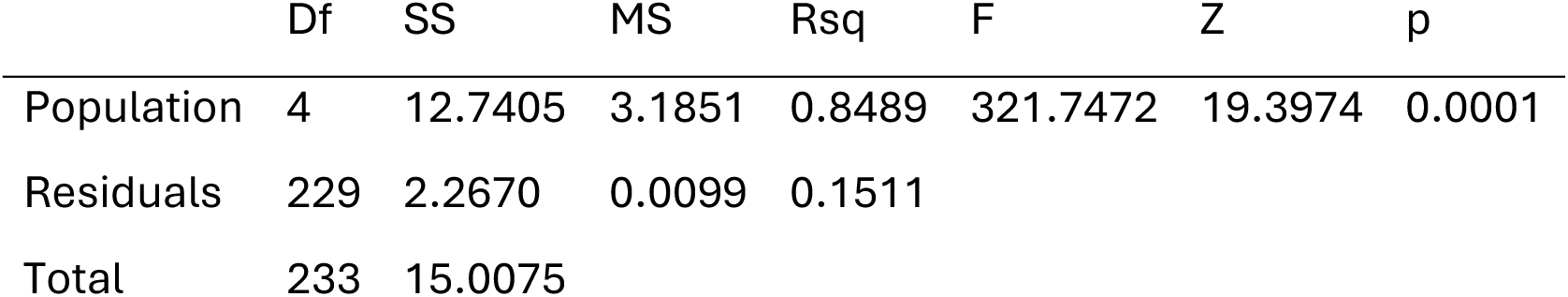
RRPP-ANOVA results comparing pelvic girdle centroid size across all populations examined. See Supplementary Table 3 for significant pairwise comparisons. Abbreviations: Df = degrees of freedom; F = F statistic; MS = mean square; p = p value (10,000 permutations); Rsq = R squared; SS = sum of squares; Z = effect size.

### Shape

PC1 for the skull represented 27.5% of the total variance and described an axis of variation that differentiated marine and freshwater phenotypes, with Sarita specimens clustering among the other freshwater specimens (Figure 3a). The positive end of PC1 contained mostly marine individuals and depicted dorsoventrally expanded skulls with a more dorsally oriented mandible, dorsally extended opercle and narrower orbits. The negative end of PC1 contained mostly freshwater individuals and depicted dorsoventrally flattened skulls with a reduced mandibular angle, shorter opercles and larger, wider orbits. Within these habitat groupings, individual populations largely overlapped without obvious differentiation along this axis. While overlapping with the other freshwater populations, a substantial portion of the Sarita Lake cluster occupied a more positive region of the PC1 axis, closer to the marine populations, than did the other freshwater populations.

**Figure 3.**
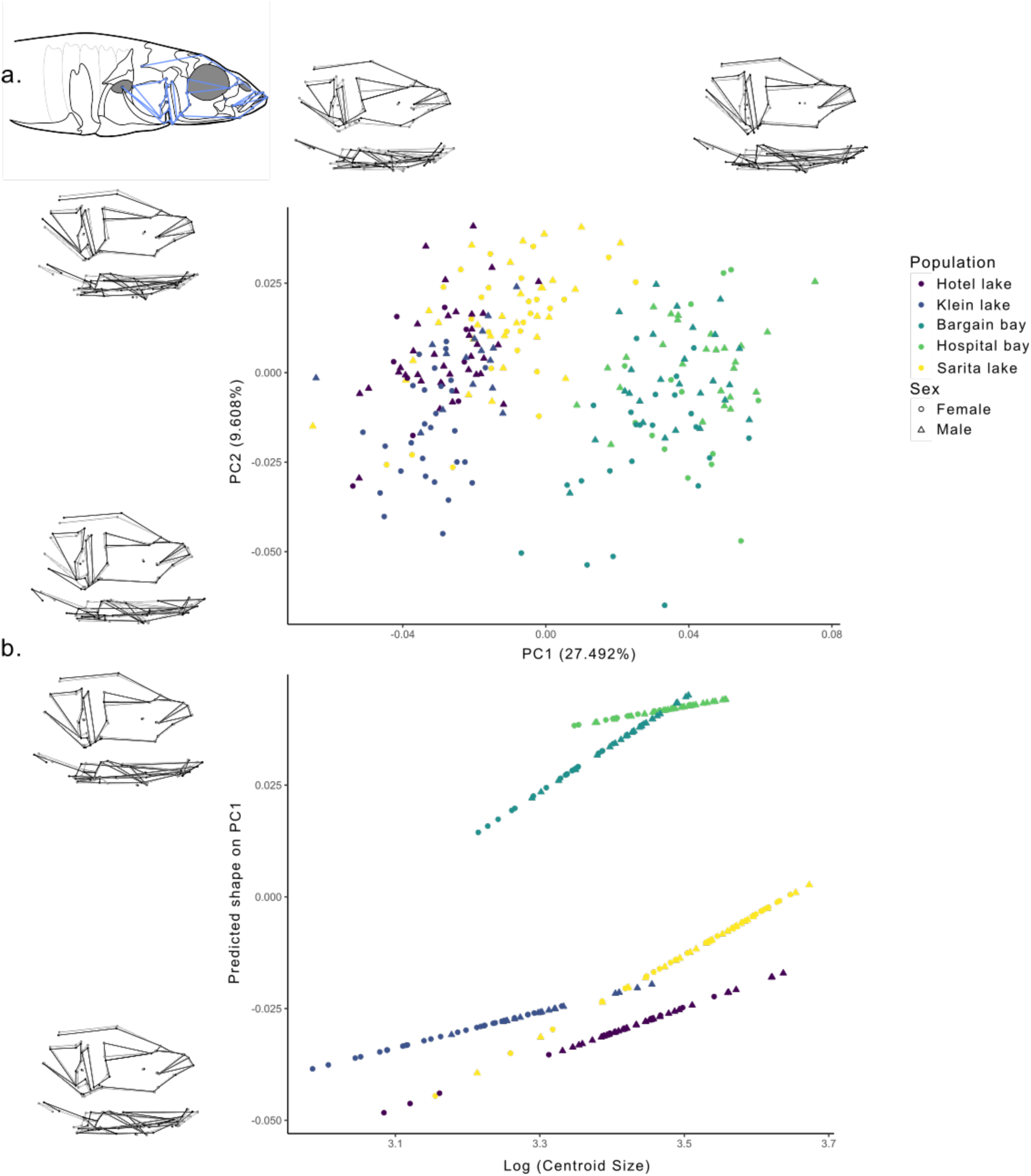
Morphological and allometric variation in the skull. a. Principal component analysis of skull shape. b. Allometric trajectories for the shape of the skull across populations and sexes. Plotted points in b. represent predicted shape scores on PC1 based on the fitted values from the model presented in Table 5. Wireframes along principal components illustrate shape deformation in the skull in lateral (top) and superior (bottom) views at the extreme ends of each component. Black lines indicate the shape at the extremes and the grey lines indicate the mean shape.

**Table 5.**
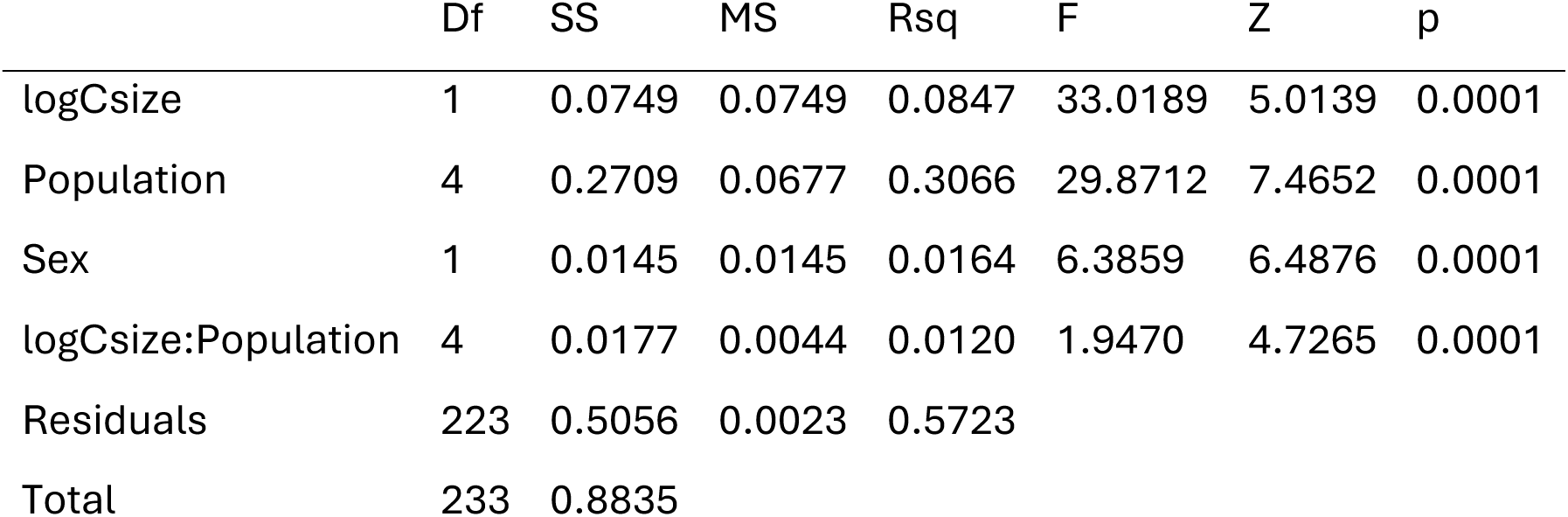
Procrustes RRPP-ANOVA results comparing skull shape across all populations examined. Abbreviations: Df = degrees of freedom; F = F statistic; Log CSize = natural log of centroid size; MS = mean square; p = p value (10,000 permutations); Rsq = R squared; SS = sum of squares; Z = effect size.

|  | Df | SS | MS | Rsq | F | Z | p |
| --- | --- | --- | --- | --- | --- | --- | --- |
| logCsize | 1 | 0.0749 | 0.0749 | 0.0847 | 33.0189 | 5.0139 | 0.0001 |
| Population | 4 | 0.2709 | 0.0677 | 0.3066 | 29.8712 | 7.4652 | 0.0001 |
| Sex | 1 | 0.0145 | 0.0145 | 0.0164 | 6.3859 | 6.4876 | 0.0001 |
| logCsize:Population | 4 | 0.0177 | 0.0044 | 0.0120 | 1.9470 | 4.7265 | 0.0001 |
| Residuals | 223 | 0.5056 | 0.0023 | 0.5723 |  |  |  |
| Total | 233 | 0.8835 |  |  |  |  |  |

PC2 for the skull represented 9.6% of the total variance and did not clearly differentiate populations. All six populations varied widely along the PC2 axis, with similar variance in both the freshwater and marine groups. The negative end of PC2 depicted individuals with dorsally expanded skulls, taller opercles, and shorter subopercles, while the positive end of PC2 contained individuals with dorsally flattened skulls, shorter opercles, and much longer subopercles. A substantial portion of the Sarita Lake cluster occupied a more positive region of the PC2 axis than either the marine or freshwater populations.

PC1 in the pectoral girdle analysis represented 38.5% of total variance and described an axis of variation that primarily differentiated the Sarita Lake population from the other four populations (Figure 4a). The positive end of PC1 depicted individuals with overall antero-posterior widening, and an ectocoracoid that extended distinctly more posterior and ventrally. The negative end of PC1 depicted individuals with overall antero-posterior shortening, and an ectocoracoid extending relatively more dorsal and anterior.

**Figure 4.**
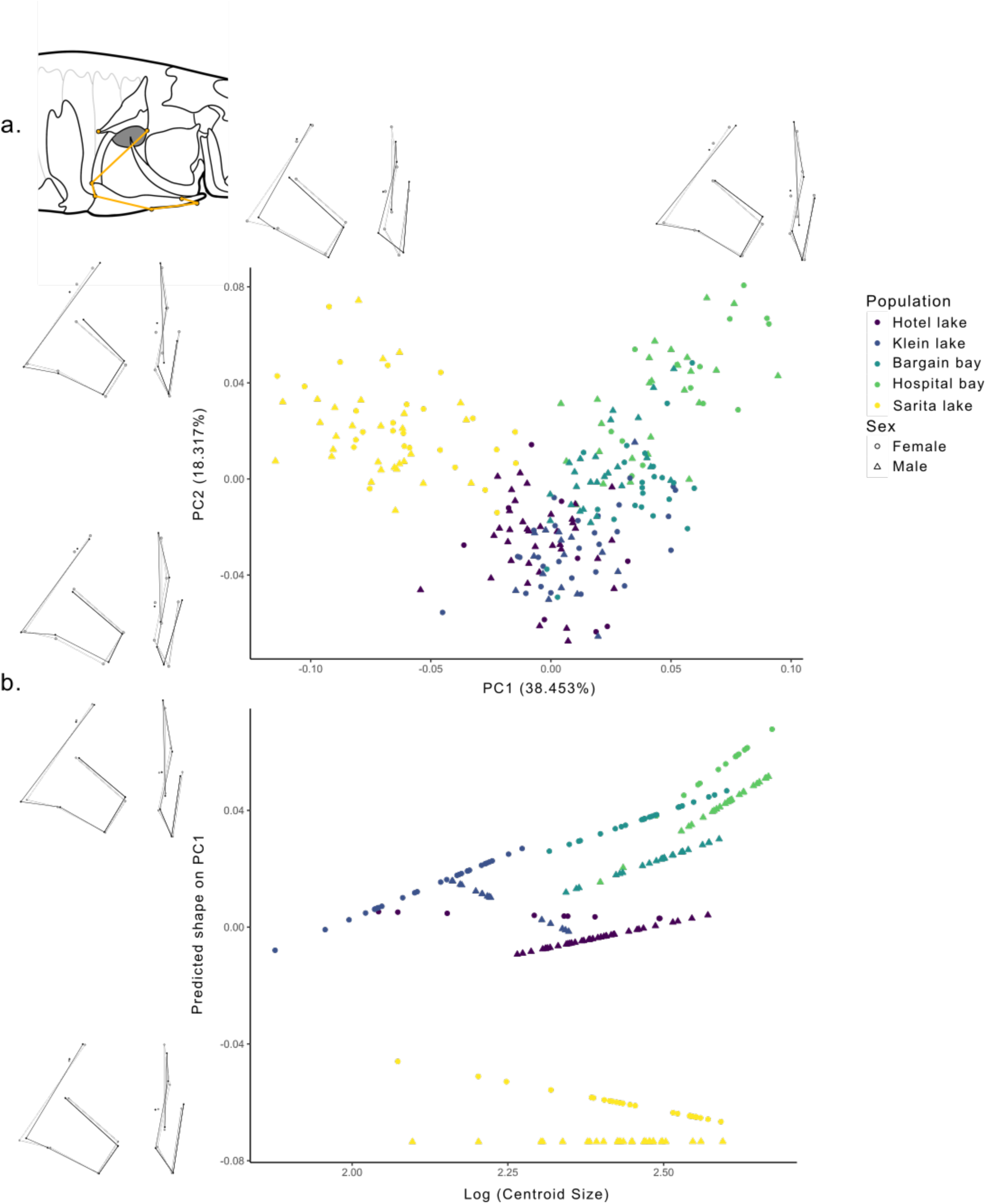
Morphological and allometric variation in the pectoral girdle. a. Principal component analysis of pectoral girdle shape. b. Allometric trajectories for the shape of the pectoral girdle across populations and sexes. Plotted points in b. represent predicted shape scores on PC1 based on the fitted values from the model presented in Table 6. Wireframes along principal components illustrate shape deformation the pectoral girdle in lateral (left) and frontal (right) views at the extreme end of each component. Black lines indicate the shape at the extremes and the grey lines indicate the mean shape.

**Table 6.** Procrustes RRPP-ANOVA results comparing pectoral girdle shape across all populations examined. Abbreviations: Df = degrees of freedom; F = F statistic; log CSize = natural log of centroid size; MS = mean square; p = p value (10,000 permutations); Rsq = R squared; SS = sum of squares; Z = effect size.

|  | Df | SS | MS | Rsq | F | Z | p |
| --- | --- | --- | --- | --- | --- | --- | --- |
| logCsize | 1 | 0.1088 | 0.1088 | 0.0872 | 40.3351 | 6.7137 | 0.001 |
| Population | 4 | 0.4833 | 0.1208 | 0.3872 | 44.7820 | 10.1584 | 0.0001 |
| Sex | 1 | 0.0300 | 0.0301 | 0.0241 | 11.1277 | 6.5871 | 0.0001 |
| logCsize:Population | 4 | 0.0173 | 0.0043 | 0.0139 | 1.6105 | 2.1589 | 0.0144 |
| logCsize:Sex | 1 | 0.0036 | 0.0036 | 0.0029 | 1.3289 | 0.8204 | 0.2096 |
| Population:Sex | 4 | 0.0120 | 0.0030 | 0.0096 | 1.1098 | 0.5408 | 0.2893 |
| logCsize:Population:Sex | 4 | 0.0156 | 0.0039 | 0.0125 | 1.4436 | 1.6976 | 0.0469 |
| Residuals | 214 | 0.5774 | 0.0027 | 0.4626 |  |  |  |
| Total | 233 | 1.2481 |  |  |  |  |  |

PC2 in the pectoral girdle analysis represented 18.3% of total variance and described an axis of variation that partially separated Klein Lake and Hotel Lake from the other three populations. The positive end of PC2 depicted individuals with overall antero-posterior shortening and dorsoventral elongation of the girdle, with a more anterior ectocoracoid that extended farther cranially. The negative end of PC2 depicted individuals with an overall antero-posterior widening and dorsoventral shortening of the girdle, with a more posterior ectocoracoid with a reduced cranial extension.

Unlike in the skull, the pectoral girdle PCA did not clearly differentiate populations along marine-freshwater lines, but rather indicated that Sarita Lake pectoral girdles occupy a unique region of the morphospace relative to the other groups. Marine forms were limited to the positive end of PC1, and mostly occupied the positive PC1 and positive PC2 quadrants. Of the marine populations, Hospital Bay extended closer to the extremes of this quadrant than Bargain Bay. There was no single freshwater shape space as Klein and Hotel Lakes, and Sarita Lake largely occupied separate quadrants. Klein Lake and Hotel Lake were primarily located at the negative end of PC2 and clustered closer to the PC1 midline, whereas Sarita Lake was uniquely located in the negative PC1 and positive PC2 quadrant.

PC1 in the pelvic girdle analysis represented 25.6% of total variance and described an axis of variation that primarily differentiated between Sarita Lake and the other four populations (Figure 5a). The positive end of PC1 depicted individuals with antero-posterior compression of the girdle, a shorter and more posterior angled ascending process, and a longer pelvic plate. The negative end of PC1 depicted individuals with some antero-posterior expansion, particularly towards the base of the ascending process, as well as an overall vertically oriented ascending process with a shorter pelvic plate. PC2 in the pelvic girdle analysis represented 19.9% of total variance, and primarily differentiated between the freshwater and marine populations. The positive end of PC2 depicted individuals with some antero-posterior expansion of the girdle, a much taller ascending process, and a slightly shorter pelvic plate. The negative end of PC2 depicted individuals with some antero-posterior compression of the girdle, a much shorter ascending process, and a longer pelvic plate. Interestingly, both principal components possess the same pattern where shorter ascending processes are paired with a longer pelvic plate, and vice versa. Similar to the results for the pectoral girdle, the Sarita Lake specimens clustered in their own quadrant of the morphospace, separate from all other populations.

**Figure 5.**
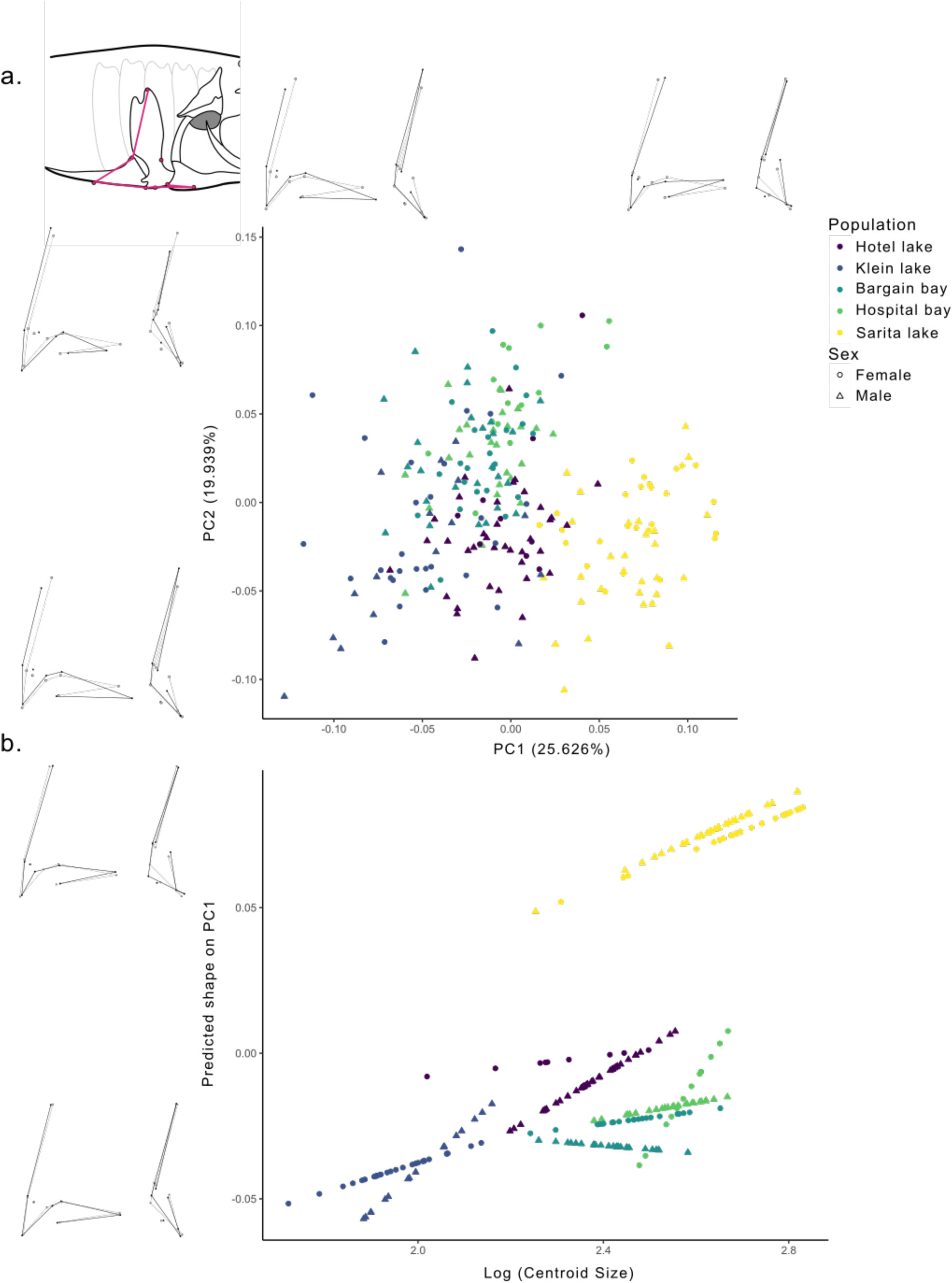
Morphological and allometric variation in the pelvic girdle. a. Principal component analysis of pelvic girdle shape. b. Allometric trajectories for the shape of the pelvic girdle across populations and sexes. Plotted points in b. represent predicted shape scores on PC1 based on the fitted values from the model presented in Table 7. Wireframes along principal components illustrate shape deformation the pelvic girdle in lateral (left) and frontal (right) views at the extreme end of each component. Black lines indicate the shape at the extremes and the grey lines indicate the mean shape.

**Table 7.** Procrustes RRPP-ANOVA results comparing pelvic girdle shape across all populations examined. Abbreviations: Df = degrees of freedom; F = F statistic; log CSize = natural log of centroid size; MS = mean square; p = p value (10,000 permutations); Rsq = R squared; SS = sum of squares; Z = effect size.

|  | Df | SS | MS | Rsq | F | Z | p |
| --- | --- | --- | --- | --- | --- | --- | --- |
| logCsize | 1 | 0.2938 | 0.2938 | 0.1363 | 51.8774 | 9.6238 | 0.0001 |
| Population | 4 | 0.4641 | 0.1160 | 0.2153 | 20.4873 | 14.0552 | 0.0001 |
| Sex | 1 | 0.0510 | 0.0510 | 0.0237 | 9.0123 | 5.4761 | 0.0001 |
| logCsize:Population | 4 | 0.0420 | 0.0105 | 0.0195 | 1.8520 | 2.4946 | 0.0061 |
| logCsize:Sex | 1 | 0.0219 | 0.0219 | 0.0102 | 3.8649 | 3.2188 | 0.0004 |
| Population:Sex | 4 | 0.0303 | 0.0076 | 0.0140 | 1.3359 | 1.2503 | 0.1051 |
| logCsize:Population:Sex | 4 | 0.0410 | 0.0103 | 0.0190 | 1.8100 | 2.3938 | 0.0085 |
| Residuals | 214 | 1.2120 | 0.0057 | 0.5621 |  |  |  |
| Total | 233 | 2.1562 |  |  |  |  |  |

### Allometry

There were significant effects of size, population, and sex, as well as a significant interaction between size and population (but not sex) on the shape of the skull (Table 5). The significant interaction between size and population suggested the presence of variation in allometric effects on skull shape among populations. Pairwise comparisons of distances between least squares means reveal significant differences in skull morphology across all populations (p < 0.05, Supplementary Table 4). Populations displayed similar levels of morphological disparity, despite Bargain Bay displaying significantly more variance than Hotel Lake and Klein Lake (Supplementary Table 5). Regressing PC1 for the fitted values against logCsize (Figure 4b) showed that the relationship between shape and size among the skulls of these populations was variable, with relatively steeper slopes in Sarita Lake and Bargain Bay populations compared to the other three populations indicating greater changes to skull shape relative to each unit of size. Taken together, these results support that differences in allometric variation are linked with differences in disparity across populations and contribute to observed shape differences. Because sex was not a significant factor in this model, we did not plot separate regressions for males and females.

There were significant effects of size, population and sex on the shape of the pectoral girdle, a significant interaction between size and population, and a significant three-way interaction among size, population and sex indicating variable allometric effects associated with sex within some populations (Table 6). Pectoral girdle morphology was significantly different between nearly all population by sex combinations, with the exception of male:female comparisons in Hospital Bay, Hotel Lake, Klein Lake and Sarita Lake (Supplementary Table 6). Hotel Lake females were also not significantly different from Klein Lake females. In contrast to what was observed for the skull, morphological disparity is nearly uniform across the pectoral girdle for all populations, with no significant comparisons across populations or sexes (Supplementary Table 7). Regressing PC1 for the fitted values against logCsize (Figure 4b) once again showed that the relationship between shape and size in the pectoral girdle of these populations is variable. Of particular note, there is substantial sexually mediated variation in this relationship for the freshwater populations, where male and female slopes are very different from each other in both magnitude and direction for all three freshwater populations, suggesting substantial allometric variation between sexes within populations as well as among populations.

There were significant effects of size, population and sex on the shape of the pelvic girdle, significant interactions between size and population and size and sex, and a significant three-way interaction among size, population and sex once again indicating variable allometric effects associated with sex within some populations (Table 7). Similar to the pectoral girdle, pelvic girdle morphology was significantly different between nearly all population by sex combinations (Supplementary Table 8). Male and female pelvic morphologies were not significantly different in Bargain Bay, Hotel Lake and Sarita Lake. Pairwise comparisons failed to indicate significant differences in pelvic morphology between Hospital Bay males and Bargain Bay males and females. Interestingly, pairwise comparisons of variance indicate that Klein Lake males and females are significantly more morphologically disparate than any of the other groups (p < 0.05, Supplementary Table 9), but these two groups did not differ in variance between themselves. Regressing PC1 for the fitted values against logCsize (Figure 5b) once again showed considerable variability across groups, but in contrast to the other body regions, here the Sarita Lake population was the only one with similar slopes for males and females. All other populations showed different relationships between shape and size for males compared to females within the same populations, with the freshwater populations showing a steeper slope for males in both cases, while the marine populations had steeper slopes for females, indicating greater changes to pelvic girdle shape relative to each unit of size in freshwater males and marine females vs their counterparts within each population.

## Discussion

We tested the hypothesis that the apparently ‘giant’ phenotype of the Sarita Lake stickleback population is produced by allometric changes in the skull, pectoral, and pelvic girdles, and that the phenotype of these individuals is otherwise similar to that of individuals from nearby freshwater populations. We found that changing relationships between size and shape are key for producing the phenotypic variation we observed in the Sarita Lake specimens, and that these relationships are different among body regions and between sexes within and across populations. We found that, rather than being uniformly larger, the Sarita Lake population is differently larger than most other populations in each body region, and that this observation is driven in part by female morphology.

### Sarita Lake Phenotype and Implications for Ecology

We found that the Sarita Lake females had relatively larger skulls and pectoral girdles than those of the other freshwater populations we examined (Figure 2) and the Sarita Lake pelvic girdles were larger than all others except Hospital Bay Lagoon (Figure 2). Notably, there was considerable size variation among all populations in both girdles that was not reflected in the skull. Taken together, these results suggest that the Sarita Lake population does tend to be larger than other nearby populations, and that this effect is largely driven by females. Sexually-mediated size variation in threespine stickleback skulls and pectoral girdles is less well understood than shape variation, which is discussed further below. In contrast, threespine stickleback are known to exhibit sexually-mediated size variation in the pelvic girdle, with the female pelvic girdle tending to be larger and longer than that of males (Kitano et al., 2007, Aguirre et al., 2008). Female-biased sexually-mediated size variation has been implicated in gigantism in ninespine stickleback (*Pungitius pungitius*), where females tend to be larger and more variable in giant populations (Herczeg et al., 2010). This trait is thought to be linked to fecundity selection as the underlying cause of gigantism, with larger females having a higher reproductive output (Herczeg et al., 2010).

Sarita Lake skulls were similar in shape overall to the skulls of the other freshwater populations, and diverged from those of the marine populations (Figure3a), although they retained some marine-like features including a relatively dorsoventrally expanded skull and relatively dorsally expanded opercle. Notably, and in contrast to our findings for size, we did not demonstrate sexually-mediated variation within populations for the skull, and once we controlled for the effects of size, we were unable to recover significant shape differences among populations but rather demonstrated very high levels of morphological disparity, suggesting that size is strongly influencing shape in the skull across populations. In contrast, the Sarita Lake girdles were unique relative to all other groups examined, with a prominent anteroposterior compression of the pectoral girdle with a more dorsal and anterior ectocoracoid, a slightly shorter pelvic ascending process, and elongated pelvic plate. Shape disparity among populations was less apparent in the girdles, and after controlling for the effects of size we did not recover any significant differences between Sarita Lake and the other populations. Notably, we observed significant sexually-mediated variation in the Klein Lake girdles after controlling for size, and further found that the Klein Lake females were significantly smaller than most other groups for these two body regions (Figure 2), effectively the opposite of what we observed for Sarita Lake, indicating an interesting direction for future study of the Klein Lake population.

Collectively, our phenotypic analysis of Sarita Lake fish describes a population with relatively dorsally expanded skulls, relatively narrowed pectoral girdles, and relatively robust pelvic armour, in which females tended to be larger than males and where size variation was heavily implicated in structuring shape variation. The opercle serves as a critical part of the threespine stickleback trophic apparatus, and as such, opercle shape and size variation are strongly associated with dietary variation (Caldecutt and Adams, 1998, Kimmel et al., 2012, McGee et al., 2013). Taller opercles like those of the Sarita Lake individuals are associated with marine and freshwater limnetic populations (Caldecutt and Adams, 1998, Taugbol et al., 2020, McGee et al., 2013), suggesting that the Sarita Lake population largely occupies a limnetic niche in this habitat. Further, gill raker number has been shown to be positively correlated with opercle size (Caldecutt and Adams, 1998), suggesting a relationship between opercle size and diet. We also note that many trophic traits are known to be plastic and capable of rapid change in threespine stickleback (Bell and Foster, 1994, Leaver and Reimchen, 2012), which may underly the high levels of morphological disparity we report in the skull. However, despite this disparity, our results are consistent with previous work demonstrating that the effect of habitat on shape is very strong in the skull (Schutz et al., 2022) with the Sarita Lake population clustering more closely with freshwater populations for this body region relative to the others, and that substantial interpopulation variation is often outweighed by macro-level habitat effects in these fish (Kimmel et al., 2012).

Threespine stickleback rely on pectoral fin rowing for locomotion (Bell and Foster, 1994), and the ecotocoracoid is an important site of muscle attachment for the pectoral fins. Most of what is known about locomotor variation in this taxon, however, is largely based on traits of the pectoral fin itself (Hermida et al., 2005), which was not part of this study. Variation in pelvic morphology of threespine stickleback, in contrast, is commonly linked with predation pressure. The pelvic girdle directly supports the pelvic spines, and the ascending process of this girdle has a key role in bracing the dorsal spines, all of which are a critical component in the threespine stickleback anti-predator apparatus (Reimchen, 1988, Bell et al., 1985, Bell et al., 1993, Miller et al., 2017). As such, an increase in pelvic plate length may suggest that the Sarita Lake population is subject to a more intense predation regime. Variation in pelvic morphology is consistently associated with variation in predation pressure in threespine stickleback, as the ascending process, plate and spines form a critical defensive complex, and pelvic reduction has been repeatedly found to be associated with reduced predation pressure (Bell et al., 1993, Miller et al., 2017, Klepaker and Østbye, 2008, Lescak and von Hippel, 2011). Conversely, higher levels of fish predation favour increased pelvic armour (Miller et al., 2017). The relatively large overall size of the pelvic girdle in the Sarita Lake population further supports the notion that predation pressure is relatively strong in this habitat. This finding is consistent with other literature on gigantism in stickleback linking unusually large body size with predation pressure from both predatory fish and gape-limited predators in general (Bell, 1984, Reimchen, 1991, Reimchen, 1988, Moodie, 1972, Moodie and Reimchen, 1976). The ‘giant’ Haida Gwaii stickleback had elongated pelvic spines, and an examination of predatory fish stomach contents in the area found a preference for smaller fish with shorter spines (Moodie, 1972). Furthermore, predator-induced injuries indicating survival of a predation event were found to be more common in larger sticklebacks (Reimchen, 1988). Future work to characterize the predation regime on threespine stickleback in Sarita Lake will help to test the hypothesis that predation regime is driving size and shape changes in these fish.

Like many similar studies, we note that the present work represents a snapshot approach to comparing morphological variation by sampling localities once per year, in this case including specimens from two consecutive years. Such a sampling approach does not account for plasticity in shape (e. g. Sharpe et al., 2008, Hoffmann and Borg, 2006, Spoljaric and Reimchen, 2011, Wund et al., 2012, McGuigan et al., 2010), indeterminate growth, seasonal and temperature effects (e. g. Lefebure et al., 2011), or other sources of variability that may not be completely captured by a restricted sampling paradigm. Future such studies would benefit from a longitudinal approach to better capture the relationship between size and shape across time in each population to more thoroughly characterize differences among them.

### Developmental and Evolutionary Integration in Sarita Lake

The allometric slopes recovered in our analysis differed widely across all groups considered and across body regions, and no consistent patterns were observed. In the skull, the Sarita Lake population had a relatively steeper positive allometric slope than did the other freshwater populations, which were nearly identical (Figure 3b), indicating a different relationship between size and shape in this body region for this population that was not influenced by sexually-mediated variation. The allometric slopes in the girdles revealed a complex relationship between size and shape, with the strongest effects of sexually-mediated variation present in the pectoral girdle of the freshwater populations (Figure 4b). The relationship between size and shape in Sarita Lake pectoral girdles was driven by females, where we observed a negative relationship associated with nearly no relationship in Sarita Lake males. Interestingly, in the pelvic girdle, the Sarita Lake population demonstrated a relatively steep positive slope and, in contrast to the other populations, nearly no sexually-mediated variation, whereas the males of the Klein and Hotel Lake populations demonstrated considerably steeper slopes than their female counterparts (Figure 5b). Taken together, these results show that there is no consistent relationship between size and shape across body regions within the Sarita Lake population altogether or even within Sarita Lake females, despite their tendency toward larger size in each region (Supplementary Tables 1, 2 and 3). These fish are therefore not isometrically larger than their nearby counterparts, but rather demonstrate a complex pattern of development that is producing larger body elements in different ways across the skeleton, effectively using different developmental pathways to reach large size in different regions. This result is consistent with those of many other studies that have shown clear links between allometric and phenotypic variation in stickleback (Aguirre and Bell, 2012, Kimmel et al., 2008, McGuigan et al., 2010, Spoljaric and Reimchen, 2011, Taugbol et al., 2020, Wund et al., 2012).The results presented here further suggest variation in the developmental integration between the skull, pectoral and pelvic girdles across populations, which may be implicated in the large size of the Sarita Lake population. Indeed, correlation and covariation patterns among body regions can be produced by many different factors, and these patterns are constantly overwritten during ontogeny at the level of individual organisms, producing a ‘palimpsest’ effect over the course of development (Hallgrimsson et al., 2009). During ontogeny, plasticity in response to environmental factors can significantly affect between-trait relationships and can both introduce and reduce covariation between traits. Predation is known to induce plasticity across stickleback populations, which may then display markedly different reaction norms (Bell et al., 1993, Miller et al., 2017, Klepaker and Østbye, 2008, Lescak and von Hippel, 2011). Recent studies of stickleback populations in Greenland show that populations co-occurring with Arctic Char are characterized by a larger body size and a larger skull size than populations exempt from predation (Moosmann et al., 2025). However, reaction norms between the two body regions are not correlated, indicating that environmental pressures such as predation regime and prey availability could have decoupled those traits.

Organismal growth over the course of ontogeny is among the strongest mechanisms responsible for phenotypic integration, acting to maintain functional relationships among anatomical components as both overall body size and local element size changes (Zelditch, 1988, Cheverud, 1996, Klingenberg, 2016, Hallgrimsson et al., 2009). Ontogenetic changes pertaining to growth rate of body depth and head length relative to standard length were observed amongst Japanese threespine stickleback populations occupying estuarine and marine habitats, further supporting a role of plasticity in shaping allometry and covariation amongst traits (Kanbe et al., 2025).

In threespine stickleback, studies have shown that genes related to the transition from marine to freshwater environments can contribute to the generation of phenotypic variation and changes in the patterning of covariation across traits. For instance, the ectodysplasin gene (*Eda*) and its receptor (*Edar*) are well-known contributors to the production of morphological variation in threespine stickleback (Colosimo et al., 2005, O’Brown et al., 2015, Laurentino et al., 2022). *Eda* has an important role in the development of bone and neural tissue (Rodriguez-Ramirez et al., 2023) and has been shown to produce pleiotropic effects across the body (Aguirre and Bell, 2012, Rogers et al., 2012). Notably, a quantitative trait locus of major effect that impacts both pelvic girdle and skull morphology has been mapped in close proximity to *Eda* (Albert et al., 2008), suggesting that this locus could modulate shape integration systems such as the skull and pelvic girdle.

We did not conduct any direct tests of relationship between genetic loci and the phenotypic variation we report here, which limits our ability to suggest causal links between developmental integration and variational modularity in the present study. Future work would benefit from approaches that directly link genotype to phenotype, such as quantitative trait locus analysis (e.g. Jamniczky et al., 2015), to more clearly identify the genetic drivers for the variation observed here.

### Conclusions

We describe a population of threespine stickleback from Sarita Lake, BC that are relatively larger than nearby populations across the skull and girdles. This increase in size may be driven by relatively increased predation pressure in Sarita Lake. Interestingly, this size variation is not uniform and rather than being isometrically larger, this population presents a complex pattern of phenotypic variation where size and shape are heavily interconnected, such that each body region is larger in a different way relative to other populations. Further, female size increase is primarily driving the observation that Sarita Lake fish are larger than others. As many others have noted, sexually mediated variation is an important contributor to the variation that we observe, and further work targeting the intersection of predation and sex-specific behaviour differences may help elucidate the mechanisms producing phenotypic variation in Sarita Lake threespine stickleback.

## Author Contributions

SP led the data collection and analysis. SP, KD and HJ analysed data. SP wrote the first draft of the manuscript. SP, KD and HJ edited the manuscript and approved the final version. HJ obtained funding to support the work.

## Data Availability

Data and code will be deposited in Dryad upon acceptance.

## Acknowledgements

The University of Calgary is situated within the traditional territories of the people of the Treaty 7 region in Southern Alberta, which includes the Blackfoot Confederacy (comprising the Siksika, Piikani, and Kainai First Nations), as well as the Tsuut’ina First Nation, and the Stoney Nakoda (including the Chiniki, Bearspaw, and Wesley First Nations). The City of Calgary is also home to Métis Nation within Alberta, including Nose Hill Metis District 5 and Elbow Metis District 6. We acknowledge and pay tribute to these peoples, along with the traditional caretakers of the lands in which we collected samples and performed our work, including the Huu-ay-aht First Nations and the Squamish, Sechelt, and Tla’amin and Klahoose nations. We would also like to acknowledge the following individuals for their support, which was critical to the completion of this project: Heidi Schutz, Sean Rogers, Kelsey Lucas, and the Jamniczky and Lucas Labs at the University of Calgary. The Natural Sciences and Engineering Research Council of Canada provided funds to HJ to support this work.

