## Supplementary Tables for "Disentangling shape and size in a population of unusually large Threespine Stickleback (*Gasterosteus aculeatus*) from Vancouver Island, British Columbia"

SUPPLEMENTARY INFORMATION

Supplementary Table 1. Pairwise comparisons of centroid size across the skull for all studied populations and sexes. D, distance; UCL (95%), upper confidence limit of 95% ; Z, Z-score. Significant p-values are indicated in bold.

| Comparison | d | UCL (95%) | Z | p-value |
|---|---|---|---|---|
| Hotel Lake Female : Klein Lake Female | 0.157548 | 0.222228 | -0.33236 | 0.632 |
| Hotel Lake Female : Bargain Bay Female | 0.007226 | 0.082719 | -1.27603 | 0.8806 |
| Hotel Lake Female : Hospital Bay Female | 0.098699 | 0.114352 | 1.242972 | 0.1048 |
| Hotel Lake Female : Sarita Lake Female | 0.182393 | 0.167658 | 2.069494 | **0.0207** |
| Hotel Lake Female : Hotel Lake Male | 0.121936 | 0.132132 | 1.344509 | 0.0871 |
| Hotel Lake Female : Klein Lake Male | 0.010444 | 0.150231 | -2.32668 | 0.989 |
| Hotel Lake Female : Bargain Bay Male | 0.09041 | 0.108779 | 1.144518 | 0.1247 |
| Hotel Lake Female : Hospital Bay Male | 0.158275 | 0.185811 | 0.881475 | 0.1869 |
| Hotel Lake Female : Sarita Lake Male | 0.189247 | 0.243288 | 0.108382 | 0.4579 |
| Klein Lake Female: Bargain Bay Female | 0.164774 | 0.183455 | 0.907332 | 0.1865 |
| Klein Lake Female : Hospital Bay Female | 0.256247 | 0.266639 | 1.292169 | 0.0977 |
| Klein Lake Female : Sarita Lake Female | 0.339941 | 0.316866 | 2.593563 | **0.005** |
| Klein Lake Female : Hotel Lake Male | 0.279484 | 0.281642 | 1.552105 | 0.0604 |
| Klein Lake Female : Klein Lake Male | 0.147104 | 0.123226 | 2.471817 | **0.0051** |
| Klein Lake Female : Bargain Bay Male | 0.247958 | 0.259579 | 1.174121 | 0.1179 |
| Klein Lake Female : Hospital Bay Male | 0.315823 | 0.335048 | 0.808358 | 0.2101 |
| Klein Lake Female : Sarita Lake Male | 0.346795 | 0.393745 | -0.31902 | 0.6265 |
| Bargain Bay Female : Hospital Bay Female | 0.091473 | 0.125544 | 0.495706 | 0.318 |
| Bargain Bay Female : Sarita Lake Female | 0.175167 | 0.177672 | 1.557886 | 0.0579 |
| Bargain Bay Female : Hotel Lake Male | 0.11471 | 0.142043 | 0.50169 | 0.3113 |
| Bargain Bay Female : Klein Lake Male | 0.01767 | 0.113534 | -1.74201 | 0.9508 |
| Bargain Bay Female : Bargain Bay Male | 0.083184 | 0.118663 | 0.228791 | 0.4115 |
| Bargain Bay Female : Hospital Bay Male | 0.15105 | 0.19642 | -0.13172 | 0.553 |
| Bargain Bay Female : Sarita Lake Male | 0.182021 | 0.253134 | -1.22137 | 0.8874 |
| Hospital Bay Female : Sarita Lake Female | 0.083694 | 0.107166 | 0.876715 | 0.1921 |
| Hospital Bay Female : Hotel Lake Male | 0.023237 | 0.072274 | -0.14056 | 0.5716 |
| Hospital Bay Female : Klein Lake Male | 0.109143 | 0.195127 | -1.00616 | 0.8412 |
| Hospital Bay Female : Bargain Bay Male | 0.008289 | 0.057422 | -0.80836 | 0.7768 |
| Hospital Bay Female : Hospital Bay Male | 0.059576 | 0.126265 | -0.60012 | 0.7272 |
| Hospital Bay Female : Sarita Lake Male | 0.090548 | 0.183753 | -1.58514 | 0.9429 |
| Sarita Lake Female : Hotel Lake Male | 0.060457 | 0.069254 | 1.241906 | 0.1056 |
| Sarita Lake Female : Klein Lake Male | 0.192837 | 0.24759 | -0.26021 | 0.6052 |
| Sarita Lake Female : Bargain Bay Male | 0.091983 | 0.098578 | 1.385556 | 0.0815 |
| Sarita Lake Female : Hospital Bay Male | 0.024117 | 0.058773 | 0.172207 | 0.4498 |
| Sarita Lake Female : Sarita Lake Male | 0.006854 | 0.116947 | -3.02722 | 0.9992 |
| Hotel Lake Male : Klein Lake Male | 0.132381 | 0.212015 | -1.34484 | 0.9122 |

| | | | | |
|---|---|---|---|---|
| Hotel Lake Male : Bargain Bay Male | 0.031526 | 0.063414 | 0.337226 | 0.3983 |
| Hotel Lake Male : Hospital Bay Male | 0.036339 | 0.089114 | -0.66619 | 0.7514 |
| Hotel Lake Male : Sarita Lake Male | 0.067311 | 0.145126 | -1.91553 | 0.9722 |
| Klein Lake Male : Bargain Bay Male | 0.100854 | 0.189052 | -1.48414 | 0.9307 |
| Klein Lake Male : Hospital Bay Male | 0.16872 | 0.26672 | -1.83975 | 0.9699 |
| Klein Lake Male : Sarita Lake Male | 0.199691 | 0.323684 | -2.96354 | 0.9986 |
| Bargain Bay Male : Hospital Bay Male | 0.067865 | 0.116982 | -0.37139 | 0.6505 |
| Bargain Bay Male : Sarita Lake Male | 0.098837 | 0.17454 | -1.51449 | 0.9346 |
| Hospital Bay Male : Sarita Lake Male | 0.030972 | 0.097628 | -1.14704 | 0.8678 |

Supplementary Table 2. Pairwise comparisons of centroid size across the pectoral girdle, for all studied populations and sexes. D, distance; UCL (95%), upper confidence limit of 95% ; Z, Z-score. Significant p-values are indicated in bold.

| Comparison | d | UCL (95%) | Z | p-value |
|---|---|---|---|---|
| Hotel Lake Female : Klein Lake Female | 0.171673 | 0.248783 | -0.62336 | 0.7343 |
| Hotel Lake Female : Bargain Bay Female | 0.176901 | 0.179506 | 1.597946 | **0.0568** |
| Hotel Lake Female : Hospital Bay Female | 0.305939 | 0.293455 | 1.94718 | **0.0259** |
| Hotel Lake Female : Sarita Lake Female | 0.155852 | 0.136128 | 2.104425 | **0.0168** |
| Hotel Lake Female : Hotel Lake Male | 0.099682 | 0.093829 | 1.702685 | **0.0363** |
| Hotel Lake Female : Klein Lake Male | 0.052887 | 0.21919 | -2.59792 | 0.995 |
| Hotel Lake Female : Bargain Bay Male | 0.199487 | 0.214859 | 1.255272 | 0.1019 |
| Hotel Lake Female : Hospital Bay Male | 0.300849 | 0.323305 | 1.040196 | 0.15 |
| Hotel Lake Female : Sarita Lake Male | 0.135297 | 0.169781 | 0.712028 | 0.2325 |
| Klein Lake Female : Bargain Bay Female | 0.348573 | 0.353906 | 1.448037 | 0.0758 |
| Klein Lake Female : Hospital Bay Female | 0.477612 | 0.469642 | 1.929964 | **0.0269** |
| Klein Lake Female : Sarita Lake Female | 0.327524 | 0.308842 | 2.394913 | **0.0081** |
| Klein Lake Female : Hotel Lake Male | 0.271354 | 0.266636 | 1.837094 | **0.0343** |
| Klein Lake Female : Klein Lake Male | 0.118786 | 0.083151 | 2.454877 | **0.0016** |
| Klein Lake Female : Bargain Bay Male | 0.37116 | 0.388618 | 0.95423 | 0.1692 |
| Klein Lake Female : Hospital Bay Male | 0.472521 | 0.497617 | 0.632433 | 0.2621 |
| Klein Lake Female : Sarita Lake Male | 0.30697 | 0.344261 | 0.138119 | 0.4371 |
| Bargain Bay Female : Hospital Bay Female | 0.129039 | 0.159907 | 0.623963 | 0.273 |
| Bargain Bay Female : Sarita Lake Female | 0.021049 | 0.086524 | -0.91131 | 0.8049 |
| Bargain Bay Female : Hotel Lake Male | 0.077219 | 0.122572 | -0.21492 | 0.5863 |
| Bargain Bay Female : Klein Lake Male | 0.229787 | 0.325258 | -1.50224 | 0.9318 |
| Bargain Bay Female : Bargain Bay Male | 0.022587 | 0.079665 | -0.57253 | 0.7123 |
| Bargain Bay Female : Hospital Bay Male | 0.123948 | 0.189735 | -0.83394 | 0.7996 |
| Bargain Bay Female : Sarita Lake Male | 0.041604 | 0.0527 | 1.169761 | 0.1252 |
| Hospital Bay Female : Sarita Lake Female | 0.150088 | 0.202278 | -0.09973 | 0.54 |
| Hospital Bay Female : Hotel Lake Male | 0.206258 | 0.238195 | 0.48229 | 0.314 |
| Hospital Bay Female : Klein Lake Male | 0.358826 | 0.440126 | -0.75204 | 0.7715 |
| Hospital Bay Female : Bargain Bay Male | 0.106452 | 0.123325 | 1.091366 | 0.1372 |
| Hospital Bay Female : Hospital Bay Male | 0.005091 | 0.087568 | -1.61913 | 0.9337 |
| Hospital Bay Female : Sarita Lake Male | 0.170642 | 0.166205 | 1.824229 | **0.0357** |
| Sarita Lake Female : Hotel Lake Male | 0.05617 | 0.077787 | 0.764288 | 0.233 |
| Sarita Lake Female : Klein Lake Male | 0.208738 | 0.280577 | -0.78448 | 0.7825 |
| Sarita Lake Female : Bargain Bay Male | 0.043636 | 0.1211 | -1.42001 | 0.9189 |
| Sarita Lake Female : Hospital Bay Male | 0.144997 | 0.229795 | -1.74357 | 0.9587 |
| Sarita Lake Female : Sarita Lake Male | 0.020555 | 0.077034 | -0.66124 | 0.7404 |
| Hotel Lake Male : Klein Lake Male | 0.152568 | 0.238878 | -1.46308 | 0.9286 |

| | | | | |
|---|---|---|---|---|
| Hotel Lake Male : Bargain Bay Male | 0.099806 | 0.157463 | -0.80681 | 0.7876 |
| Hotel Lake Male : Hospital Bay Male | 0.201167 | 0.266507 | -1.1437 | 0.8733 |
| Hotel Lake Male : Sarita Lake Male | 0.035615 | 0.112635 | -1.72398 | 0.9565 |
| Klein Lake Male : Bargain Bay Male | 0.252374 | 0.359563 | -2.02524 | 0.979 |
| Klein Lake Male : Hospital Bay Male | 0.353735 | 0.469572 | -2.37609 | 0.9914 |
| Klein Lake Male : Sarita Lake Male | 0.188184 | 0.315533 | -2.91581 | 0.9984 |
| Bargain Bay Male : Hospital Bay Male | 0.101361 | 0.150753 | -0.32366 | 0.6283 |
| Bargain Bay Male : Sarita Lake Male | 0.06419 | 0.084667 | 0.845529 | 0.2064 |
| Hospital Bay Male : Sarita Lake Male | 0.165552 | 0.193682 | 0.495886 | 0.3125 |

Supplementary Table 3. Pairwise comparisons of centroid size across the pelvic girdle for all studied populations. D, distance; UCL (95%), upper confidence limit of 95% ; Z, Z-score. Significant p-values are indicated in bold.

| Comparison | D | UCL (95%) | Z | p-value |
|---|---|---|---|---|
| Hotel Lake : Klein Lake | 0.386732 | 0.105967 | 4.735294 | **1.00E-04** |
| Hotel Lake : Bargain Bay | 0.082402 | 0.103175 | 1.209679 | 0.1166 |
| Hotel Lake : Hospital Bay | 0.19499 | 0.110256 | 2.746344 | **5.00E-04** |
| Hotel Lake : Sarita Lake | 0.296511 | 0.100326 | 4.03946 | **1.00E-04** |
| Klein Lake : Bargain Bay | 0.469134 | 0.103743 | 5.503704 | **1.00E-04** |
| Klein Lake : Hospital Bay | 0.581722 | 0.110219 | 5.980304 | **1.00E-04** |
| Klein Lake : Sarita Lake | 0.683243 | 0.102025 | 6.774208 | **1.00E-04** |
| Bargain Bay : Hospital Bay | 0.112588 | 0.106181 | 1.697328 | **0.0374** |
| Bargain Bay : Sarita Lake | 0.214109 | 0.097227 | 3.295128 | **1.00E-04** |
| Hospital Bay : Sarita Lake | 0.101521 | 0.10503 | 1.538791 | 0.0586 |

Supplementary Table 4. Pairwise comparisons of shape across the skull for all studied populations. Populations are compared, and sex is excluded, following the results of the reduced linear model for the skull. D, distance; UCL (95%), upper confidence limit of 95% ; Z, Z-score. Significant p-values are indicated in bold.

| Comparison | d | UCL (95%) | Z | p-value |
|---|---|---|---|---|
| Bargain Bay : Hospital Bay | 0.023007 | 0.017858 | 2.568446 | **0.0043** |
| Bargain Bay : Hotel Lake | 0.066623 | 0.017203 | 5.217139 | **1.00E-04** |
| Bargain Bay : Klein Lake | 0.063867 | 0.01751 | 4.977814 | **1.00E-04** |
| Bargain Bay : SaritaLake | 0.056476 | 0.016435 | 4.828586 | **1.00E-04** |
| Hospital Bay : HotelLake | 0.073088 | 0.018304 | 5.197961 | **1.00E-04** |
| Hospital Bay : KleinLake | 0.073849 | 0.018559 | 5.040925 | **1.00E-04** |
| Hospital Bay : SaritaLake | 0.063057 | 0.017362 | 4.999679 | **1.00E-04** |
| Hotel Lake : KleinLake | 0.034291 | 0.01762 | 3.784291 | **1.00E-04** |
| Hotel Lake : SaritaLake | 0.04681 | 0.016777 | 4.435133 | **1.00E-04** |
| Klein Lake : SaritaLake | 0.046336 | 0.01701 | 4.436412 | **1.00E-04** |

Supplementary Table 5. Pairwise comparisons of variance across the skull for all studied populations. Populations are compared, and sex is excluded, following the results of the reduced linear model for the skull. D, distance; UCL (95%), upper confidence limit of 95% ; Z, Z-score. Significant p-values are indicated in bold.

| Comparison | d | UCL (95%) | Z | P-value |
|---|---|---|---|---|
| Bargain Bay : Hospital Bay | 0.000174 | 0.000567 | -0.09585 | 0.5499 |
| Bargain Bay : Hotel Lake | 0.000649 | 0.000538 | 1.937904 | **0.0196** |
| Bargain Bay : Klein Lake | 0.000647 | 0.000539 | 1.92274 | **0.0182** |
| Bargain Bay : Sarita Lake | 0.000288 | 0.000524 | 0.635595 | 0.2804 |
| Hospital Bay : Hotel Lake | 0.000475 | 0.000579 | 1.25905 | 0.1041 |
| Hospital Bay : Klein Lake | 0.000472 | 0.000578 | 1.227956 | 0.1097 |
| Hospital Bay : Sarita Lake | 0.000114 | 0.000555 | -0.51967 | 0.6938 |
| Hotel Lake : Klein Lake | 2.25E-06 | 0.000547 | -2.29969 | 0.995 |
| Hotel Lake : Sarita Lake | 0.000361 | 0.000533 | 0.943044 | 0.1832 |
| Klein Lake : Sarita Lake | 0.000359 | 0.000548 | 0.916408 | 0.189 |

Supplementary Table 6. Pairwise comparisons of shape across the pectoral girdle for all studied populations and sexes. D, distance; UCL (95%), upper confidence limit of 95% ; Z, Z-score. Significant p-values are indicated in bold.

| Comparison | d | UCL (95%) | Z | p-value |
|---|---|---|---|---|
| Bargain Bay Female : Hospital Bay Female | 0.055916 | 0.037093 | 3.120979 | **0.0009** |
| Bargain Bay Female : Hotel Lake Female | 0.057711 | 0.042855 | 2.705202 | **0.0023** |
| Bargain Bay Female : Klein Lake Female | 0.056154 | 0.030595 | 3.809436 | **1.00E-04** |
| Bargain Bay Female : Sarita Lake Female | 0.10301 | 0.031397 | 5.87995 | **1.00E-04** |
| Bargain Bay Female : Bargain Bay Male | 0.035593 | 0.031456 | 2.093827 | **0.0177** |
| Bargain Bay Female : Hospital Bay Male | 0.058579 | 0.031378 | 3.869315 | **1.00E-04** |
| Bargain Bay Female : Hotel Lake Male | 0.063206 | 0.02873 | 4.560881 | **1.00E-04** |
| Bargain Bay Female : Klein Lake Male | 0.069646 | 0.034979 | 4.044316 | **1.00E-04** |
| Bargain Bay Female : Sarita Lake Male | 0.116035 | 0.030298 | 6.629423 | **1.00E-04** |
| Hospital Bay Female : Hotel Lake Female | 0.093527 | 0.046729 | 4.138103 | **1.00E-04** |
| Hospital Bay Female : Klein Lake Female | 0.084741 | 0.035788 | 4.673457 | **1.00E-04** |
| Hospital Bay Female : Sarita Lake Female | 0.11958 | 0.036826 | 5.606683 | **1.00E-04** |
| Hospital Bay Female : Bargain Bay Male | 0.057204 | 0.036207 | 3.293133 | **0.0004** |
| Hospital Bay Female : Hospital Bay Male | 0.031983 | 0.036079 | 1.23896 | 0.1103 |
| Hospital Bay Female : Hotel Lake Male | 0.093609 | 0.034342 | 5.263303 | **1.00E-04** |
| Hospital Bay Female : Klein Lake Male | 0.095935 | 0.039724 | 4.788698 | **1.00E-04** |
| Hospital Bay Female : Sarita Lake Male | 0.132888 | 0.035662 | 6.153284 | **1.00E-04** |
| Hotel Lake Female : Klein Lake Female | 0.029344 | 0.041358 | 0.458724 | 0.3228 |
| Hotel Lake Female : Sarita Lake Female | 0.086869 | 0.041848 | 4.336095 | **1.00E-04** |

| | | | | |
|---|---|---|---|---|
| Hotel Lake Female : Bargain Bay Male | 0.057048 | 0.041925 | 2.726641 | **0.0026** |
| Hotel Lake Female : Hospital Bay Male | 0.082786 | 0.041933 | 4.092428 | **1.00E-04** |
| Hotel Lake Female : Hotel Lake Male | 0.031275 | 0.040185 | 0.772352 | 0.2215 |
| Hotel Lake Female : Klein Lake Male | 0.04612 | 0.04507 | 1.737102 | **0.0416** |
| Hotel Lake Female : Sarita Lake Male | 0.094386 | 0.041635 | 4.596848 | **1.00E-04** |
| Klein Lake Female : Sarita Lake Female | 0.09025 | 0.029552 | 5.715502 | **1.00E-04** |
| Klein Lake Female : Bargain Bay Male | 0.046893 | 0.029986 | 3.279107 | **0.0004** |
| Klein Lake Female : Hospital Bay Male | 0.070994 | 0.029823 | 4.584497 | **1.00E-04** |
| Klein Lake Female : Hotel Lake Male | 0.029868 | 0.027224 | 1.97117 | **0.0252** |
| Klein Lake Female : Klein Lake Male | 0.025675 | 0.034092 | 0.680486 | 0.2516 |
| Klein Lake Female : Sarita Lake Male | 0.099217 | 0.02897 | 6.016457 | **1.00E-04** |
| Sarita Lake Female : Bargain Bay Male | 0.087152 | 0.030492 | 5.342078 | **1.00E-04** |
| Sarita Lake Female : Hospital Bay Male | 0.106856 | 0.030615 | 6.006979 | **1.00E-04** |
| Sarita Lake Female : Hotel Lake Male | 0.077039 | 0.028235 | 4.988836 | **1.00E-04** |
| Sarita Lake Female : Klein Lake Male | 0.092598 | 0.034606 | 5.223003 | **1.00E-04** |
| Sarita Lake Female : Sarita Lake Male | 0.019489 | 0.029629 | 0.234299 | 0.4027 |
| Bargain Bay Male : Hospital Bay Male | 0.042235 | 0.030356 | 2.75873 | **0.0031** |
| Bargain Bay Male : Hotel Lake Male | 0.049452 | 0.027909 | 3.711392 | **1.00E-04** |
| Bargain Bay Male : Klein Lake Male | 0.051563 | 0.034676 | 3.019726 | **0.0009** |
| Bargain Bay Male : Sarita Lake Male | 0.098344 | 0.029555 | 5.900927 | **1.00E-04** |
| Hospital Bay Male : Hotel Lake Male | 0.07593 | 0.027948 | 5.074348 | **1.00E-04** |
| Hospital Bay Male : Klein Lake Male | 0.077331 | 0.034505 | 4.48622 | **1.00E-04** |
| Hospital Bay Male : Sarita Lake Male | 0.117511 | 0.029641 | 6.41611 | **1.00E-04** |
| Hotel Lake Male : Klein Lake Male | 0.033692 | 0.032221 | 1.813874 | **0.0337** |
| Hotel Lake Male : Sarita Lake Male | 0.082359 | 0.027065 | 5.660593 | **1.00E-04** |
| Klein Lake Male : Sarita Lake Male | 0.099481 | 0.033671 | 5.564864 | **1.00E-04** |

Supplementary Table 7. Pairwise comparisons of variance across the pectoral girdle for all studied populations and sexes. D, distance; UCL (95%), upper confidence limit of 95% ; Z, Z-score. Significant p-values are indicated in bold.

| Comparison | d | UCL (95%) | Z | P-value |
|---|---|---|---|---|
| Bargain Bay Female : Hospital Bay Female | 0.000558 | 0.000985 | 0.739243 | 0.2379 |
| Bargain Bay Female : Hotel Lake Female | 0.000765 | 0.001126 | 0.983788 | 0.1653 |
| Bargain Bay Female : Klein Lake Female | 0.000835 | 0.000808 | 1.655501 | 0.0427 |
| Bargain Bay Female : Sarita Lake Female | 0.000385 | 0.000805 | 0.456772 | 0.342 |
| Bargain Bay Female : Bargain Bay Male | 0.000136 | 0.000827 | -0.6423 | 0.7327 |
| Bargain Bay Female : Hospital Bay Male | 0.000195 | 0.000805 | -0.32703 | 0.6278 |
| Bargain Bay Female : Hotel Lake Male | 2.82E-05 | 0.000757 | -1.6357 | 0.9412 |
| Bargain Bay Female : Klein Lake Male | 3.38E-05 | 0.00096 | -1.61313 | 0.9388 |
| Bargain Bay Female : Sarita Lake Male | 0.000113 | 0.000798 | -0.79357 | 0.775 |
| Hospital Bay Female : Hotel Lake Female | 0.000207 | 0.001238 | -0.60324 | 0.7241 |

| | | | | |
|---|---|---|---|---|
| Hospital Bay Female : Klein Lake Female | 0.000277 | 0.000937 | -0.07394 | 0.5385 |
| Hospital Bay Female : Sarita Lake Female | 0.000173 | 0.000946 | -0.54352 | 0.7014 |
| Hospital Bay Female : Bargain Bay Male | 0.000694 | 0.000965 | 1.072361 | 0.1478 |
| Hospital Bay Female : Hospital Bay Male | 0.000363 | 0.000951 | 0.215677 | 0.431 |
| Hospital Bay Female : Hotel Lake Male | 0.000586 | 0.000891 | 0.918516 | 0.1855 |
| Hospital Bay Female : Klein Lake Male | 0.000524 | 0.001075 | 0.533731 | 0.3105 |
| Hospital Bay Female : Sarita Lake Male | 0.000445 | 0.000939 | 0.474377 | 0.3351 |
| Hotel Lake Female : Klein Lake Female | 6.98E-05 | 0.001093 | -1.31399 | 0.8937 |
| Hotel Lake Female : Sarita Lake Female | 0.00038 | 0.001121 | 0.105523 | 0.4734 |
| Hotel Lake Female : Bargain Bay Male | 0.000901 | 0.001134 | 1.241327 | 0.1043 |
| Hotel Lake Female : Hospital Bay Male | 0.00057 | 0.001122 | 0.587377 | 0.2886 |
| Hotel Lake Female : Hotel Lake Male | 0.000793 | 0.001063 | 1.129091 | 0.1276 |
| Hotel Lake Female : Klein Lake Male | 0.000731 | 0.001228 | 0.820993 | 0.2126 |
| Hotel Lake Female : Sarita Lake Male | 0.000652 | 0.001094 | 0.806307 | 0.2154 |
| Klein Lake Female : Sarita Lake Female | 0.00045 | 0.000782 | 0.729319 | 0.2515 |
| Klein Lake Female : Bargain Bay Male | 0.000971 | 0.000797 | 1.961339 | 0.0167 |
| Klein Lake Female : Hospital Bay Male | 0.00064 | 0.000772 | 1.266689 | 0.1033 |
| Klein Lake Female : Hotel Lake Male | 0.000863 | 0.00071 | 1.967808 | 0.0173 |
| Klein Lake Female : Klein Lake Male | 0.000801 | 0.000915 | 1.371184 | 0.082 |
| Klein Lake Female : Sarita Lake Male | 0.000722 | 0.000755 | 1.516397 | 0.06 |
| Sarita Lake Female : Bargain Bay Male | 0.000521 | 0.000804 | 0.916004 | 0.1898 |
| Sarita Lake Female : Hospital Bay Male | 0.00019 | 0.000777 | -0.31732 | 0.6256 |
| Sarita Lake Female : Hotel Lake Male | 0.000413 | 0.000719 | 0.706345 | 0.2569 |
| Sarita Lake Female : Klein Lake Male | 0.000351 | 0.000915 | 0.221633 | 0.4312 |
| Sarita Lake Female : Sarita Lake Male | 0.000272 | 0.000771 | 0.105128 | 0.4711 |
| Bargain Bay Male : Hospital Bay Male | 0.000331 | 0.000803 | 0.281598 | 0.404 |
| Bargain Bay Male : Hotel Lake Male | 0.000108 | 0.000748 | -0.78286 | 0.7716 |
| Bargain Bay Male : Klein Lake Male | 0.00017 | 0.00094 | -0.56551 | 0.7123 |
| Bargain Bay Male : Sarita Lake Male | 0.000249 | 0.000782 | -0.02878 | 0.5249 |
| Hospital Bay Male : Hotel Lake Male | 0.000223 | 0.000743 | -0.08564 | 0.5433 |
| Hospital Bay Male : Klein Lake Male | 0.000161 | 0.000911 | -0.59617 | 0.7159 |
| Hospital Bay Male : Sarita Lake Male | 8.19E-05 | 0.000778 | -1.0191 | 0.8343 |
| Hotel Lake Male : Klein Lake Male | 6.21E-05 | 0.000862 | -1.2691 | 0.8858 |
| Hotel Lake Male : Sarita Lake Male | 0.000141 | 0.000707 | -0.52148 | 0.6924 |
| Klein Lake Male : Sarita Lake Male | 7.91E-05 | 0.000912 | -1.14044 | 0.8594 |

Supplementary table 8. Pairwise comparisons of shape across the pelvic girdle for all studied populations and sexes. D, distance; UCL (95%), upper confidence limit of 95% ; Z, Z-score. Significant p-values are indicated in bold.

| Comparison | d | UCL (95%) | Z | p-value |
|---|---|---|---|---|
| Bargain Bay Female : Hospital Bay Female | 0.044982 | 0.046582 | 1.517852 | **0.0651** |
| Bargain Bay Female : Hotel Lake Female | 0.055872 | 0.053488 | 1.826179 | **0.0333** |
| Bargain Bay Female : Klein Lake Female | 0.062469 | 0.038487 | 3.87775 | **1.00E-04** |
| Bargain Bay Female : Sarita Lake Female | 0.101757 | 0.03911 | 6.28476 | **1.00E-04** |
| Bargain Bay Female : Bargain Bay Male | 0.036163 | 0.039242 | 1.318007 | 0.0963 |
| Bargain Bay Female : Hospital Bay Male | 0.033987 | 0.039072 | 1.075868 | 0.1445 |
| Bargain Bay Female : Hotel Lake Male | 0.065254 | 0.036091 | 4.38866 | **1.00E-04** |
| Bargain Bay Female : Klein Lake Male | 0.079384 | 0.044116 | 4.219689 | **1.00E-04** |
| Bargain Bay Female : Sarita Lake Male | 0.109235 | 0.037519 | 6.826641 | **1.00E-04** |
| Hospital Bay Female : Hotel Lake Female | 0.073384 | 0.05897 | 2.572608 | **0.0039** |
| Hospital Bay Female : Klein Lake Female | 0.098057 | 0.044781 | 5.154345 | **1.00E-04** |
| Hospital Bay Female : Sarita Lake Female | 0.104906 | 0.04575 | 5.338041 | **1.00E-04** |
| Hospital Bay Female : Bargain Bay Male | 0.055137 | 0.045336 | 2.492689 | **0.0059** |
| Hospital Bay Female : Hospital Bay Male | 0.049427 | 0.044873 | 2.071688 | **0.02** |
| Hospital Bay Female : Hotel Lake Male | 0.090927 | 0.042992 | 5.037651 | **1.00E-04** |
| Hospital Bay Female : Klein Lake Male | 0.113229 | 0.049978 | 5.360635 | **1.00E-04** |
| Hospital Bay Female : Sarita Lake Male | 0.120895 | 0.044766 | 6.187576 | **1.00E-04** |
| Hotel Lake Female : Klein Lake Female | 0.080896 | 0.052397 | 3.535073 | **0.0003** |
| Hotel Lake Female : Sarita Lake Female | 0.093006 | 0.052902 | 4.059232 | **1.00E-04** |
| Hotel Lake Female : Bargain Bay Male | 0.064454 | 0.052833 | 2.483968 | **0.0058** |
| Hotel Lake Female : Hospital Bay Male | 0.065499 | 0.05293 | 2.540811 | **0.005** |
| Hotel Lake Female : Hotel Lake Male | 0.041402 | 0.050945 | 0.823881 | 0.2079 |
| Hotel Lake Female : Klein Lake Male | 0.086478 | 0.057313 | 3.441681 | **0.0004** |
| Hotel Lake Female : Sarita Lake Male | 0.095389 | 0.051938 | 4.351439 | **1.00E-04** |
| Klein Lake Female : Sarita Lake Female | 0.124443 | 0.037118 | 7.372053 | **1.00E-04** |
| Klein Lake Female : Bargain Bay Male | 0.073429 | 0.03709 | 4.950929 | **1.00E-04** |
| Klein Lake Female : Hospital Bay Male | 0.079653 | 0.037205 | 5.238552 | **1.00E-04** |
| Klein Lake Female : Hotel Lake Male | 0.085663 | 0.034 | 6.020984 | **1.00E-04** |
| Klein Lake Female : Klein Lake Male | 0.047015 | 0.043058 | 2.033521 | **0.021** |
| Klein Lake Female : Sarita Lake Male | 0.12378 | 0.036215 | 7.623519 | **1.00E-04** |
| Sarita Lake Female : Bargain Bay Male | 0.113318 | 0.037951 | 6.683678 | **1.00E-04** |
| Sarita Lake Female : Hospital Bay Male | 0.102509 | 0.038171 | 6.246925 | **1.00E-04** |
| Sarita Lake Female : Hotel Lake Male | 0.105204 | 0.03507 | 6.739962 | **1.00E-04** |
| Sarita Lake Female : Klein Lake Male | 0.129393 | 0.043754 | 6.574571 | **1.00E-04** |
| Sarita Lake Female : Sarita Lake Male | 0.034933 | 0.036702 | 1.427747 | 0.0764 |
| Bargain Bay Male : Hospital Bay Male | 0.021333 | 0.037933 | -0.62216 | 0.733 |

| | | | | |
|---|---|---|---|---|
| Bargain Bay Male : Hotel Lake Male | 0.063276 | 0.034771 | 4.356588 | **1.00E-04** |
| Bargain Bay Male : Klein Lake Male | 0.076109 | 0.04344 | 4.14573 | **1.00E-04** |
| Bargain Bay Male : Sarita Lake Male | 0.114858 | 0.037227 | 7.078926 | **1.00E-04** |
| Hospital Bay Male : Hotel Lake Male | 0.068926 | 0.034937 | 4.730836 | **1.00E-04** |
| Hospital Bay Male : Klein Lake Male | 0.088353 | 0.043263 | 4.990681 | **1.00E-04** |
| Hospital Bay Male : Sarita Lake Male | 0.106267 | 0.037032 | 6.662795 | **1.00E-04** |
| Hotel Lake Male : Klein Lake Male | 0.079161 | 0.040957 | 4.613564 | **1.00E-04** |
| Hotel Lake Male : Sarita Lake Male | 0.099467 | 0.033869 | 6.774683 | **1.00E-04** |
| Klein Lake Male : Sarita Lake Male | 0.122407 | 0.043108 | 6.375425 | **1.00E-04** |

Supplementary table 9. Pairwise comparisons of variance across the pelvic girdle for all studied populations and sexes. D, distance; UCL (95%), upper confidence limit of 95% ; Z, Z-score. Significant p-values are indicated in bold.

| Comparison | d | UCL (95%) | Z | p-value |
|---|---|---|---|---|
| Bargain Bay Female : Hospital Bay Female | 0.001266 | 0.003763 | 0.08875 | 0.4744 |
| Bargain Bay Female : Hotel Lake Female | 0.003659 | 0.004404 | 1.334484 | 0.0881 |
| Bargain Bay Female : Klein Lake Female | 0.00796 | 0.003105 | 3.573625 | **1.00E-04** |
| Bargain Bay Female : Sarita Lake Female | 0.000913 | 0.003177 | -0.10346 | 0.5515 |
| Bargain Bay Female : Bargain Bay Male | 0.00234 | 0.003117 | 1.114221 | 0.1393 |
| Bargain Bay Female : Hospital Bay Male | 0.001494 | 0.003159 | 0.472075 | 0.3371 |
| Bargain Bay Female : Hotel Lake Male | 0.001169 | 0.002939 | 0.249182 | 0.4189 |
| Bargain Bay Female : Klein Lake Male | 0.007745 | 0.003629 | 3.118364 | **0.0002** |
| Bargain Bay Female : Sarita Lake Male | 0.001792 | 0.003104 | 0.754515 | 0.2426 |
| Hospital Bay Female : Hotel Lake Female | 0.002394 | 0.00488 | 0.623918 | 0.2752 |
| Hospital Bay Female : Klein Lake Female | 0.006694 | 0.003725 | 2.747674 | **0.0006** |
| Hospital Bay Female : Sarita Lake Female | 0.000352 | 0.003758 | -1.0082 | 0.8327 |
| Hospital Bay Female : Bargain Bay Male | 0.001075 | 0.003738 | -0.06419 | 0.5323 |
| Hospital Bay Female : Hospital Bay Male | 0.000229 | 0.003688 | -1.30718 | 0.8928 |
| Hospital Bay Female : Hotel Lake Male | 9.68E-05 | 0.003457 | -1.73472 | 0.954 |
| Hospital Bay Female : Klein Lake Male | 0.006479 | 0.004158 | 2.421839 | **0.0031** |
| Hospital Bay Female : Sarita Lake Male | 0.000526 | 0.003659 | -0.73678 | 0.7631 |
| Hotel Lake Female : Klein Lake Female | 0.0043 | 0.004243 | 1.639908 | **0.0473** |
| Hotel Lake Female : Sarita Lake Female | 0.002746 | 0.004265 | 0.972651 | 0.1628 |
| Hotel Lake Female : Bargain Bay Male | 0.001319 | 0.004279 | 0.056858 | 0.4888 |
| Hotel Lake Female : Hospital Bay Male | 0.002165 | 0.0043 | 0.629638 | 0.2744 |
| Hotel Lake Female : Hotel Lake Male | 0.00249 | 0.004002 | 0.889072 | 0.1803 |
| Hotel Lake Female : Klein Lake Male | 0.004086 | 0.004722 | 1.415252 | 0.0797 |
| Hotel Lake Female : Sarita Lake Male | 0.001868 | 0.004169 | 0.471147 | 0.3304 |
| Klein Lake Female : Sarita Lake Female | 0.007046 | 0.002969 | 3.414921 | **1.00E-04** |
| Klein Lake Female : Bargain Bay Male | 0.005619 | 0.00294 | 2.904663 | **0.0003** |
| Klein Lake Female : Hospital Bay Male | 0.006465 | 0.003002 | 3.179215 | **1.00E-04** |
| Klein Lake Female : Hotel Lake Male | 0.006791 | 0.002722 | 3.552773 | **1.00E-04** |
| Klein Lake Female : Klein Lake Male | 0.000215 | 0.00349 | -1.32276 | 0.8966 |
| Klein Lake Female : Sarita Lake Male | 0.006168 | 0.00288 | 3.166043 | **1.00E-04** |
| Sarita Lake Female : Bargain Bay Male | 0.001427 | 0.002993 | 0.462578 | 0.3438 |
| Sarita Lake Female : Hospital Bay Male | 0.000581 | 0.003039 | -0.50419 | 0.6909 |
| Sarita Lake Female : Hotel Lake Male | 0.000256 | 0.00273 | -1.10919 | 0.8496 |
| Sarita Lake Female : Klein Lake Male | 0.006832 | 0.003582 | 2.849905 | **0.0006** |
| Sarita Lake Female : Sarita Lake Male | 0.000878 | 0.002991 | -0.0862 | 0.5425 |
| Bargain Bay Male : Hospital Bay Male | 0.000846 | 0.003034 | -0.15887 | 0.576 |

| | | | | |
|---|---|---|---|---|
| Bargain Bay Male : Hotel Lake Male | 0.001171 | 0.002768 | 0.296782 | 0.4009 |
| Bargain Bay Male : Klein Lake Male | 0.005405 | 0.003562 | 2.38786 | **0.0051** |
| Bargain Bay Male : Sarita Lake Male | 0.000549 | 0.002954 | -0.54692 | 0.702 |
| Hospital Bay Male : Hotel Lake Male | 0.000326 | 0.002773 | -0.95985 | 0.8161 |
| Hospital Bay Male : Klein Lake Male | 0.00625 | 0.003522 | 2.693327 | **0.0012** |
| Hospital Bay Male : Sarita Lake Male | 0.000297 | 0.002975 | -1.04135 | 0.837 |
| Hotel Lake Male : Klein Lake Male | 0.006576 | 0.003317 | 2.938203 | 0.0008 |
| Hotel Lake Male : Sarita Lake Male | 0.000623 | 0.002714 | -0.38241 | 0.6477 |
| Klein Lake Male : Sarita Lake Male | 0.005953 | 0.003457 | 2.627914 | **0.002** |
